# Rpl40/eL40 ribosomal protein paralogs couple cytosolic translation to mitochondrial proteome and lipid homeostasis

**DOI:** 10.64898/2026.06.13.732051

**Authors:** Kamila P. Liput, Karolina Gościńska, Monika Stasiak, Mariusz Radkiewicz, Somayeh Shahmoradi Ghahe, Katarzyna Jonak, Róża Kucharczyk, Aneta Więsyk, Eliza Molestak, Marek Tchórzewski, Matylda Macias, Aleksandra Szybińska, Ulrike Topf

## Abstract

Ribosomal protein paralogs are increasingly implicated in the regulation of cellular metabolism and mitochondrial function. However, the mechanisms linking paralog composition of ribosomes to mitochondrial physiology remain largely unclear. Here, we investigate the two Rpl40 paralogs in the budding yeast *Saccharomyces cerevisiae* and find that deletion of either paralog induces compensatory upregulation of the remaining gene and causes mild mitochondrial stress. Despite this shared phenotype, the mutants display distinct mitochondrial adaptations. Loss of Rpl40a is accompanied by increased abundance of mitochondrial proteins, including MICOS components, whereas loss of Rpl40b leads to reduced levels of mitochondrial inner membrane proteins, including the translocase Tim22 and carrier proteins, together with increased sensitivity to membrane stress. Notably, the two mutants show opposing changes in triglyceride abundance, pointing to paralog-specific control of lipid metabolic remodeling during mitochondrial stress. These findings suggest that Rpl40 paralogs differentially modulate cellular adaptation to mitochondrial stress, linking ribosome composition to mitochondrial proteostasis and lipid homeostasis.

## INTRODUCTION

Paralogous pairs of ribosomal proteins (RPs) with high sequence similarity or identity are common in eukaryotic genomes and have been proposed to contribute to ribosome heterogeneity. Individual paralogs can have different effects on cellular processes, including actin organization, bud site selection, mRNA localization, and mitochondrial function ^1^. Differences in paralog expression, especially at the post-transcriptional level, can cause changes in paralog protein abundance and incorporation into ribosomes. Often, the major paralog is expressed under unstressed conditions, while in stress conditions, including translation inhibition by hygromycin or osmotic stress, expression of the major paralog is repressed, and the minor one is a more abundant component of the ribosome ^2,3^. The expression of paralogous ribosomal proteins is regulated by non-coding sequences, introns, and untranslated regions (UTRs), which regulate splicing efficiency and translation initiation ^4^. The effect of ribosomal paralogs on cellular metabolism is also related to distinct translatomes produced by ribosomes containing only one ribosomal protein paralog promoting translation of a specific subset of mRNAs and frequently manifests as drug resistance ^5–9^. These observations support the emerging concept that variation in ribosomal protein composition can modulate ribosome function and contribute to regulatory specialization within the translational machinery ^10^.

The formation of the mitochondrial proteome depends largely on nuclear-encoded genes whose transcripts are translated by the ribosomes in the cytosol before the encoded proteins are imported into mitochondria. Several studies have linked specific ribosomal protein paralogs to mitochondrial function in yeast ^11,12^. These observations raise key questions of how changes in ribosome composition may shape the mitochondrial proteome and enable adaptive responses to changes in cellular metabolism.

In budding yeast *Saccharomyces cerevisiae*, fermentation is the main energetic-related-pathway in glucose-containing media, even in the presence of oxygen (the Crabtree effect) ^13^. During the shift from glucose-rich to non-fermentable carbon sources, cells activate mitochondrial biogenesis, increase respiratory capacity, and remodel metabolic pathways to support oxidative phosphorylation. These adaptations require extensive changes in gene expression and protein synthesis, including the production and import of numerous mitochondrial proteins. Recent structural and proteomic analyses indicate that ribosomes themselves can be remodeled during metabolic transitions ^14^. For example, time-resolved cryo-electron microscopy studies of ribosomes from *S. cerevisiae* revealed changes in ribosomal protein composition when cells are shifted from glucose to glycerol conditions, suggesting that ribosome composition can dynamically adjust to metabolic demands ^15^. Such remodeling may contribute to adaptive translation programs required for metabolic reprogramming and mitochondrial function.

Here, we investigated the role of the paralogous genes *RPL40A* and *RPL40B* under respiratory conditions. The two paralogs encode proteins with identical amino acid sequences. Rpl40/eL40 is located on the large ribosomal subunit near the GTPase-associated center, in the vicinity of the sarcin-ricin loop and elongation-factor binding site. It is incorporated into pre-60S particles at a late stage of ribosome assembly, where it supports cytoplasmic maturation of the large ribosomal subunit ^16,17^. Functional studies demonstrated that Rpl40 contributes to cellular adaptation to mitochondrial protein import stress in yeast ^18^ and overexpression of the mammalian ortholog of eL40, UBA52, prevents mitochondrial dysfunctions in neuronal cells ^19,20^. Moreover, eL40 is required for efficient replication of certain viruses ^21,22^ and has been shown to have an extraribosomal function in DNA damage response pathway ^23^. These findings support the idea that this ribosomal protein can participate in regulatory pathways that extend beyond its canonical role within the ribosome.

In this study, we demonstrate that although deleting either *RPL40* gene results in signs of mitochondrial stress, the two mutants exhibit distinct alterations in mitochondrial protein composition, stress sensitivity, and lipid profiles. These results suggest that the abundance of Rpl40 paralogs contribute differently to mitochondrial function and to the maintenance of membrane-associated metabolism in the yeast *S. cerevisiae*. Thus, variation in paralogous ribosomal protein abundance may influence cellular responses to mitochondrial perturbation.

## RESULTS

### Deletion of paralogous genes encoding *RPL40* impairs global translation under respiratory growth conditions

We used single chromosomal deletions of *RPL40A* and *RPL40B* genes to analyse the potential impact on cytosolic translation. First, we analysed the effect of paralog deletions on global translation by radioactive labelling of newly synthesized proteins (**Figure 1A, B**). Cells were grown under fermentative (glucose) or respiratory (glycerol) conditions. While no significant difference was observed when cells grew on glucose [(**Figure 1A**), ^24^], production of newly synthesized proteins decreased under respiratory conditions, albeit more pronounced in *rpl40b*Δ deletion strain (**Figure 1B**). We separated total cell lysates of wild-type and deletion cells on a linear sucrose gradient to analyse the amount and distribution of ribosome subunits, monosome and polysomes fractions (**Supplementary Figure 1A, B**). Growth on glucose caused a mild increase in the polysome-to-monosome ratio in the *RPL40A* deletion strain compared to the wild-type cells. While growth on glycerol generally results in lower ribosome occupancy on mRNA, shown by decreased polysome peaks, the analysis did not show a significant difference between the distributions of the ribosomes on mRNA in the deletions compared to wild-type cells. We reasoned that ribosome biogenesis and translation in the yeast deletion strains might not be generally impaired, as we do not see formation of halfmers (43S intermediates in translation initiation) or accumulation of 40S or 60S subunits, but lower translational output under respiratory conditions. When cell growth was challenged by cycloheximide (CHX) treatment, *rpl40a*Δ showed a pronounced sensitivity compared to wild-type cells, pointing to translation elongation impairment, considering synergistic effect, as the CHX inhibits elongating ribosomes (**Supplementary Figure 1C**). Furthermore, deletion strains showed slower growth compared to the wild-type cells under respiratory growth, using glycerol as non-fermentable carbon source (**Figure 1C**). In order to analyse whether insufficiency of Rpl40 might cause impairment with translation under respiratory growth conditions, we characterized consequences of single paralog deletions on *RPL40* transcript (**Figure 1D**) and Rpl40 protein levels (**Figure 1E**). We designed primers that allow for specific amplification of *RPL40A* or *RPL40B* transcripts (**Supplementary Table 5**). We found that deletion of either *RPL40* paralogous gene resulted in the two-fold upregulation of the transcript of the remaining *RPL40* gene, and consequently, Rpl40 protein levels tend to increase in the single deletion strains compared to wild-type cells. Other tested ribosomal protein levels appeared unchanged compared to the wild-type strain (**Figure 1E**). Thus, our results point to the fact that there is compensatory effect during respiratory growth, which is independent of overall Rpl40 levels.

**Figure 1.**
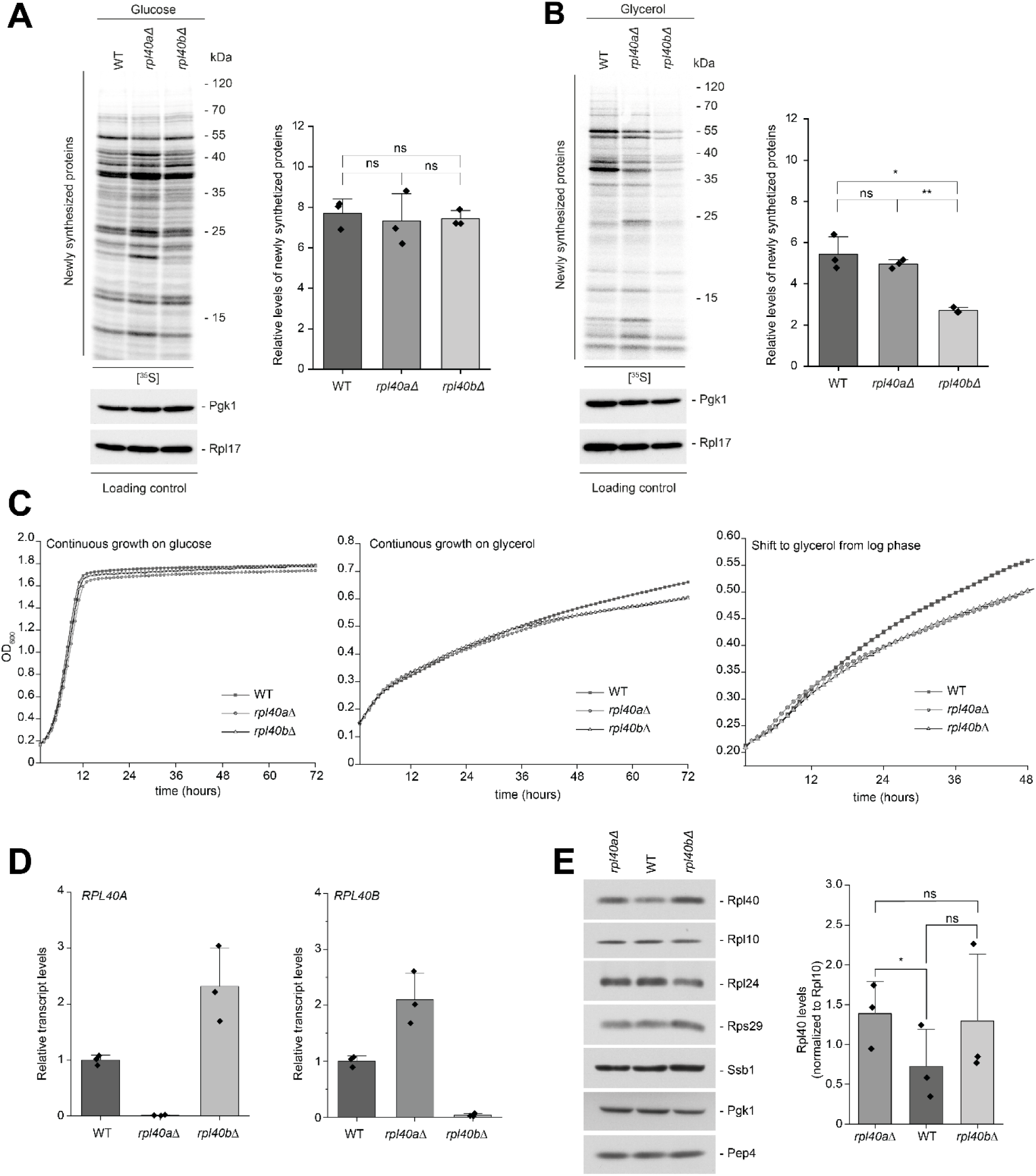
Imbalance in *RPL40* paralog abundance causes translation attenuation under respiratory conditions. A, **B.** Newly synthetized proteins of *RPL40* deletions and wild-type (WT) strain grown on glucose- (A) or glycerol-containing (B) medium. Total protein extracts were separated by SDS-PAGE and analysed by autoradiography or western blot using specific antibodies. Quantification of newly synthesized proteins relative to Rpl17 levels is presented as mean + SD. *n* = 3. Paired *t*-test was used for statistical analysis; \**p* ≤ 0.05, \*\**p* ≤ 0.01, ns (not significant) *p* > 0.05; *n* = 3. **C.** Growth curves of *RPL40* deletions and wild-type (WT) strains grown overnight in glucose-containing medium were used to start cultures in glucose-(*Left panel*) or glycerol-containing medium (*Middle panel*), respectively. Data are presented as mean of four biological replicates in three technical replicates. (*Right panel*) Cells grown in glucose-containing medium until logarithmic phase at 28°C were washed with sterile water and resuspended in glycerol-containing medium. Data are presented as mean of two biological replicates in three technical replicates. For all growth conditions, cells were incubated at 28°C and absorbance at 600 nm was measured at 60-minute intervals. **D.** Relative transcript levels of wild-type (WT), *rpl40a*Δ, and *rpl40b*Δ strains grown at 28°C on glycerol-containing medium. Transcript levels of *RPL40A* and *RPL40B* genes were calculated relative to the geometric mean of the *ACT1*, *ALG9,* and *TDH1*. Each measurement was normalized to the average gene expression of the WT strain. Data are presented as mean + SD, *n* = 3. **E.** Cells were grown on glycerol-containing medium until logarithmic phase. (*Left panel*) Total protein extracts were separated by SDS-PAGE and analysed by western blot using specific antibodies. (*Right panel*) Quantification of Rpl40 protein levels relative to Rpl10 levels. Data are presented as mean + SD. One-tailed paired *t*-test was used for statistical analysis; \**p* ≤ 0.05, ns *p* > 0.05; *n* = 3.

### Loss of Rpl40 paralogs results in mild mitochondrial stress

While nuclear genes encode the vast majority of mitochondrial proteins, some proteins governing the respiratory chain complexes are encoded by the mitochondrial genome. The cytosolic and mitochondrial translation machineries are interconnected, and changes in the cellular metabolism results in adjustment of both translation systems. We analysed the mitochondrial translation and found that while *rpl40a*Δ showed overall lower translation compared to wild-type cells, production of mitochondrial-encoded proteins was increased in *rpl40b*Δ (**Figure 2A**). Thus, deletions of the two *RPL40* genes reveal opposing modes of cytosolic-mitochondrial translational control. Mitochondrial translation also depends on the amount of mitochondrial DNA (mtDNA). Analysis of mitochondrial-encoded genes (*COX2, COX3*) in comparison to nuclear-encoded genes (*ACT1, MIP1, MRX6*) showed lower mtDNA copy number in *rpl40b*Δ, which might be compensated by the cell with increased mitochondrial mRNA translation (**Figure 2B**).

**Figure 2.**
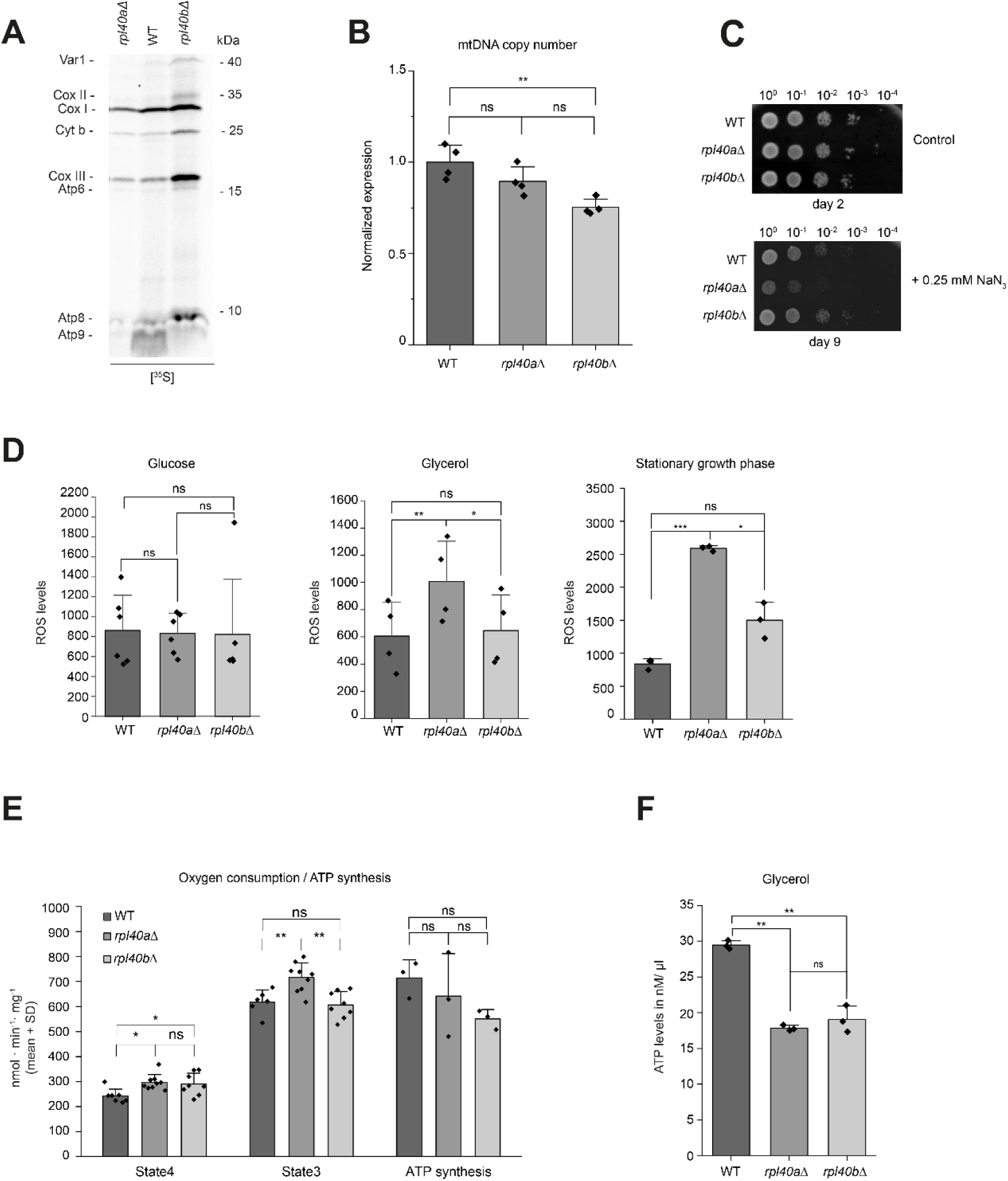
Imbalance in *Rpl40* paralog levels results in mild mitochondrial dysfunction. **A.** Mitochondrial translation assay of *RPL40* deletions and wild-type (WT) strains. Cells were grown on glycerol-containing medium. Cycloheximide was added to stop cytosolic translation. Total protein extracts were separated by SDS-PAGE and analysed by autoradiography. **B.** Relative transcript levels of selected mitochondrial-encoded genes *COX1* and *COX2* normalized to levels of nuclear-encoded genes (*ACT1*, *MIP1*, *MRX6)*. Cells grown in glycerol-containing medium until logarithmic growth phase were collected and genomic DNA was isolated. Data are presented as mean + SD. Each biological replicate was analysed in technical duplicates. Significance was determined by one-way ANOVA and Tukey test; \*\**p* ≤ 0.01, ns *p* > 0.05; *n* = 4. **C.** Growth assay of *RPL40* deletions and wild-type (WT) strains grown on glucose-containing solid plates in the presence or absence of 0.25 mM sodium azide (NaN_3_). 10-fold serial dilutions were spotted onto plates, which were then incubated at 25°C for two and nine days, respectively. Representative image of four independent experiments is shown. **D.** Measurement of reactive oxygen species (ROS) production in *RPL40* deletions and wild-type (WT) cells grown on full medium containing glucose (*left; n* = 6) or glycerol (*middle; n* = 4) to logarithmic growth phase, or in minimal medium containing glucose to stationary phase (48 hours of incubation; *right; n* = 3). Cells were washed, incubated with dihydroethidium and absorbance was measured at 5-minute intervals for 2 hours. Data are presented as mean + SD of the slope value. Significance was determined using two-tailed paired *t*-test; ns *p* > 0.05, \**p* ≤ 0.05, \*\**p* ≤ 0.01, \*\*\**p* ≤ 0.001. **E.** Measurement of oxygen consumption and ATP synthesis rates in *RPL40* deletions and wild-type (WT) strains. Measurements were performed on freshly isolated mitochondria. The state 4 and state 3 were measured when NADH and NADH/ADP were added, respectively. Oxygen consumption was quantified in nmol O₂·min⁻¹·mg⁻¹, and ATP synthesis was measured in nmol ATP·min⁻¹·mg⁻¹. Data are presented as mean + SD; *n* = 3. Significance was determined using one-way ANOVA; \**p ≤* 0.05, \*\**p* ≤ 0.01, ns *p* > 0.05. **F**. Measurement of total ATP levels in *RPL40* deletions and wild-type (WT) strains. Cells were grown on glycerol-containing medium to logarithmic growth phase. Twenty OD_600_ units of yeast cells were lysed and total ATP levels were measured using luciferase-based assay. Data are presented as mean + SD.; *n* = 3. Significance was determined using one-tailed paired *t*-test; \*\**p* ≤ 0.01, ns *p* > 0.05.

To further evaluate mitochondrial function, we analysed the growth of wild-type and deletion cells treated with sodium azide (NaN3). Sodium azide binds to the heme a3 iron and CuB sites of cytochrome c oxidase, the last enzyme complex in the electron transport chain. Thus, it blocks the electrons from being transferred to oxygen, resulting in an increased electron leakage on complex III. Strikingly, we observed a substantial decrease of growth of *rpl40a*Δ strain compared to wild-type and *rpl40b*Δ cells grown in glucose-containing medium supplemented with NaN3 (**Figure 2C**). This result points to a pre-existing compromise in electron transport, which makes the cells more sensitive to the drug. Indeed, measurement of reactive oxygen species (ROS) using dihydroethidium, an indicator of superoxide formation, confirms increased ROS levels in *rpl40a*Δ when grown in glycerol but not in glucose, at the same time indicating mitochondrial metabolic perturbations in the strain lacking *RPL40A* gene (**Figure 2D** *left and middle panel, respectively*). We also observed a significant increase in ROS levels in *rpl40aΔ* cells and a tendency toward increased ROS accumulation in *rpl40bΔ* cells upon entry into the stationary growth phase, a condition associated with a metabolic shift towards mitochondrial respiration (**Figure 2D** *right panel*). Subsequently, viability tests were performed when the cells were challenged with a toxic dose of hydrogen peroxide or an apoptosis-inducing concentration of acetic acid (100 mM). The *rpl40a*Δ cells were more susceptible to these treatments than wild-type and *rpl40b*Δ cells, possibly due to pre-existing higher cellular stress (**Supplementary Figure 2A**). Furthermore, *rpl40a*Δ exhibited reduced formation of respiratory-deficient petite colonies compared to wild-type and *rpl40b*Δ cells (**Supplementary Figure 2D**). This phenotype may arise from either reduced fitness of respiratory-deficient cells under conditions dependent on respiratory competence or improved maintenance of functional mitochondrial genomes (ρ⁺ state).

To assess respiratory activity in the *RPL40* deletion strains, mitochondria were isolated from cells grown in glycerol-rich medium, and oxygen consumption was measured using NADH as an electron donor under basal (state 4) and ADP-stimulated (state 3, phosphorylating) conditions, together with ATP synthesis rates. Mitochondria from both *rpl40aΔ* and *rpl40bΔ* cells exhibited ∼20% increase in oxygen consumption in state 4; however, only *rpl40aΔ* resulted in a 16% increase in state 3 respiration, whereas *rpl40bΔ* had no effect. Notably, ATP synthesis rates did not correlate with state 3 oxygen consumption. While mitochondrial ATP production tended to decrease in *rpl40bΔ* cells, no significant effect was observed in *rpl40aΔ* cells (**Figure 2E**). However, total cellular ATP levels were significantly lower in both deletions compared to wild-type cells, suggesting that an energy-demanding cellular process uses energy produced in mitochondria (**Figure 2F**). Although oxidative phosphorylation measurements in isolated mitochondria revealed only modest changes in respiratory states and ATP production capacity, these data contrast with the reduced cellular ATP levels, elevated ROS, and increased drug sensitivity observed particularly in *rpl40a*Δ cells. These results suggest that although the mitochondrial bioenergetic capacity per organelle is largely preserved, mitochondrial function is limited at the cellular level. Consistent with this interpretation, fluorescence imaging of mitochondria-targeted GFP revealed a reduction in mitochondrial network branching in both deletion strains, indicating altered mitochondrial dynamics and decreased network connectivity **(Supplementary Figure 2B)**. Moreover, transmission electron microscopy showed a significant reduction in total mitochondrial surface area per cell, while cristae architecture remained largely unchanged (**Supplementary Figure 2C**).

Our findings point to a primary reduction in mitochondrial abundance in the *RPL40* deletion cells, rather than the induction of major ultrastructural defects in individual mitochondria. Such remodeling is a characteristic of a cellular adaptation to impaired coordination between mitochondrial-nuclear gene expression and supports the presence of a mild, chronic mitochondrial stress state in the deletion strains ^25,26^.

### Rpl40 paralogs differentially affect proteome composition

To understand the role of Rpl40 paralogs for the composition of the mitochondrial proteome, we performed label-free quantitative (LFQ) mass spectrometry of isolated mitochondria of wild-type and *RPL40* deletion cells in four biological repetitions. In total, we quantified 976 proteins common in wild-type and *RPL40* deletion strains (**Supplementary Table 1**). The proteomic signatures of the *RPL40* deletion strains appeared to be different from each other and the wild-type strain (**Supplementary Figure 3A**). Out of all identified proteins, 548 localized in the mitochondria, and a smaller fraction of proteins localized to the endoplasmic reticulum (ER; 74), nucleus (51), or other localization (113) (**Supplementary Figure 3B**). While a comparable number of proteins were significantly decreased (28 in *rpl40a*Δ and 43 in *rpl40b*Δ) or increased (69 in *rpl40a*Δ and 55 in *rpl40b*Δ) in the deletion cells compared to wild-type cells (**Figure 3A, B**), there was unexpectedly no correlation between *rpl40a*Δ and *rpl40b*Δ cells in terms of log2 fold changes of individual proteins (**Figure 3C**). To gain further insights into the differentially regulated proteins, we analysed the subcellular localization of significantly changed proteins (*p* < 0.05) and took into consideration mitochondrial sub-compartments, other co-isolated organelles and organelle contact sites, and distinguished between membrane-associated and soluble proteins (**Figure 3D**). As expected, membrane-associated proteins clustered in outer and inner membranes of mitochondria, but also the ER, whereas soluble proteins were found in the mitochondrial matrix and cytosol. Interestingly, mitochondrial inner membrane and matrix proteins were rather increased in *rpl40a*Δ but decreased in *rpl40b*Δ cells. Furthermore, *rpl40b*Δ strain showed an enrichment in significantly decreased group of proteins localizing to the ER. Given that proteins associated with membranes exhibit opposing abundance in the two *RPL40* deletion strains, we manually checked differences in proteins associated with ER membrane composition, lipid synthesis, and membrane protein quality control (**Figure 3E**). Although none of these proteins are mitochondrial, mitochondria depend on these processes because they import most lipids from the ER, the import of proteins into the mitochondrial inner membrane is highly lipid-sensitive, and mitochondrial membrane proteins require precise membrane physical properties ^27,28^. The analysis highlights several ER- and membrane-homeostasis factors that are differentially regulated between the two *RPL40* deletion strains. Multiple proteins linked to ER network maintenance (Rtn2, Pom33), glycosylation/GPI-anchor synthesis (Alg1, Ost6, Rot2), and vesicle-mediated trafficking (Sec17, Yif1) show a significant decrease in *rpl40b*Δ but not in *rpl40a*Δ cells. Proteins mediating ER-mitochondria contacts (Mdm10) and lipid transfer (Vps13) show decreased abundance in *RPL40* deletions. On the other hand, proteins important for fatty acid activation (Faa1, Faa3) and glycerolipid synthesis (Ayr1) are induced in *rpl40b*Δ cells. These partially opposing changes in key regulators of lipid handling and membrane trafficking suggest that deletion of either *RPL40* paralog differentially influences ER-dependent membrane biogenesis pathways that might have consequences for mitochondrial membrane composition and mitochondrial biogenesis.

**Figure 3.**
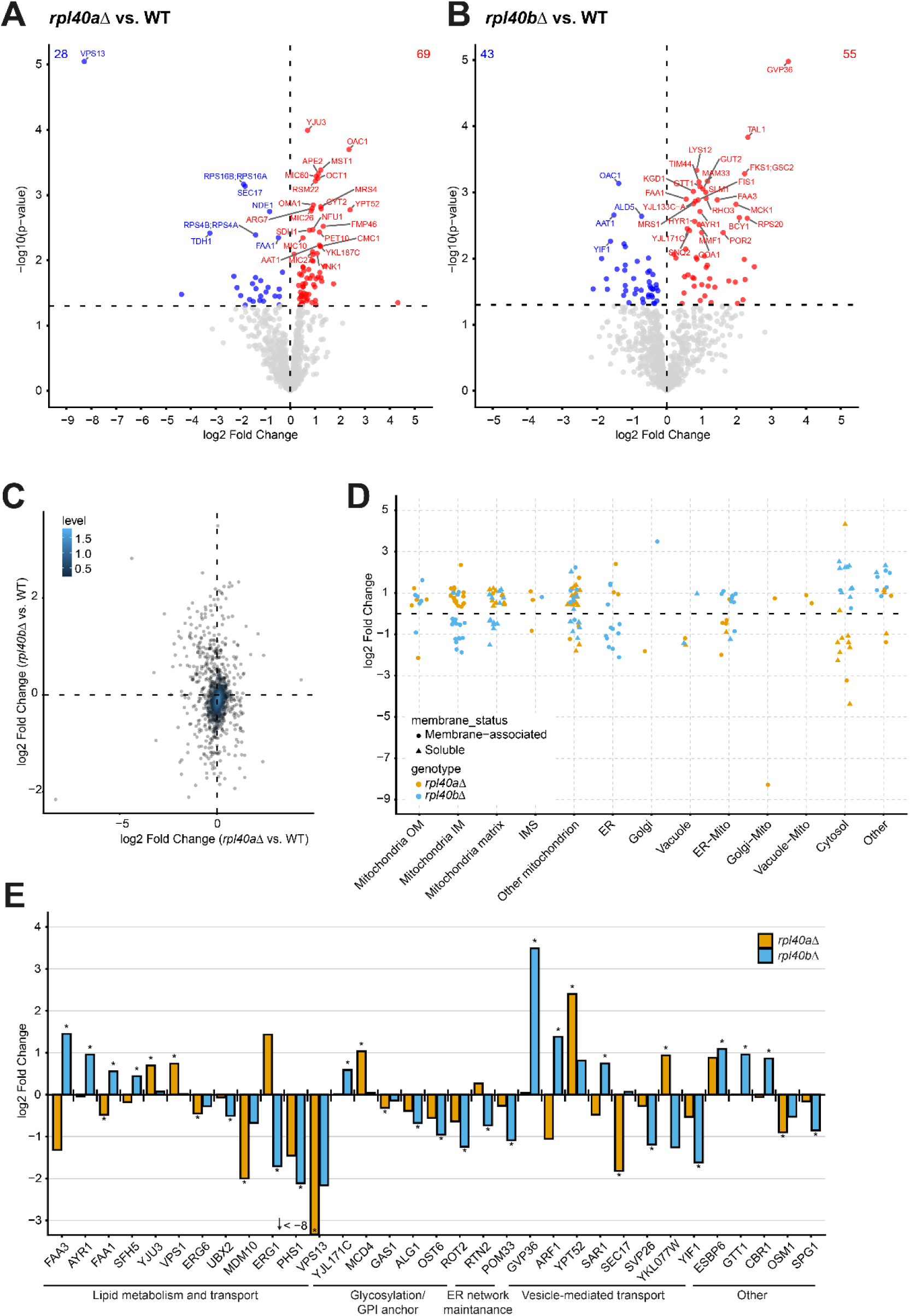
Rpl40 paralogs differentially affect proteome composition. **A, B.** Volcano plots of protein abundance changes of all quantified proteins between *rpl40a*Δ vs. wild-type (WT; A) and *rpl40b*Δ vs. WT (B) strains. Significantly regulated proteins (*p* < 0.05) are highlighted in blue for decreased abundance or in red for increased abundance. The top 20 proteins with the lowest *p*-value are labelled with the gene name. Significance was determined by Student’s *t*-test; *n* = 4. **C.** Scatter plot of log_2_ fold changes in *rpl40a*Δ versus *rpl40b*Δ strains compared to the wild type (WT). Shaded contours indicate the two-dimensional density of data points calculated by kernel density estimation. Density levels reflect relative local concentrations of data points and are not associated with statistical significance thresholds. Dashed lines mark zero-fold change. **D.** Proteins showing significant regulation (*p* < 0.05) in either *rpl40a*Δ or *rpl40b*Δ mutants were plotted according to their annotated subcellular localization and log_2_ fold change relative to wild-type cells. Cellular locations were assigned based on Gene Ontology Cellular Component (GOCC) annotations and include mitochondrial sub-compartments [outer membrane (OM), inner membrane (IM), matrix, and intermembrane space (IMS)], endoplasmic reticulum (ER), Golgi apparatus, vacuole, cytosol, ER–mitochondrial outer membrane contact sites (ER–Mito), Golgi-mitochondrial contact sites (Golgi-Mito), and vacuole and mitochondrial contact site (Vacuole-Mito). Marker color indicates genotype (*rpl40a*Δ or *rpl40b*Δ), and marker shape distinguishes membrane-associated from soluble proteins. Markers are jittered horizontally to improve visibility. The dashed horizontal line indicates no difference in fold change. **E.** Log_2_ fold changes of selected proteins are shown for *rpl40a*Δ and *rpl40b*Δ relative to wild-type (WT) cells. Proteins are grouped manually into five functional subgroups (lipid metabolism and transport; glycosylation/GPI anchor synthesis; ER network maintenance; vesicle-mediated transport; and other functions) and ordered within each subgroup by decreasing log_2_ fold change in *rpl40b*Δ. Bar colors indicate genotype (*rpl40a*Δ or *rpl40b*Δ). Asterisks denote statistically significant changes (*p* < 0.05) for the respective genotype. The horizontal line at log_2_ fold change = 0 indicates no change relative to WT.

### Rpl40 paralogs mutants show differences in membrane biogenesis or integrity

Our proteomics analysis revealed that *rpl40b*Δ strain had lower protein levels involved in lipid metabolic processes. Thus, we analysed whether the deletion mutants have a more general impact on plasma membrane composition. Sensitivity to plasma membrane damage and cell wall integrity stress is induced by sodium dodecyl sulfate (SDS) ^29^. Upon exposure to SDS, *rpl40b*Δ cells showed increased sensitivity compared to wild-type and *rpl40a*Δ cells (**Figure 4A**). No difference in growth was observed when cells were exposed to substances inducing osmotic stress, 0.5 M NaCl and 0.5 M sorbitol, which substantiates the conclusion that loss of *RPL40B* is correlated primarily with impaired cell wall and/or membrane integrity (**Figure 4B**). SDS-sensitivity could also be linked to reduced Ca^2+^-dependent plasma membrane repair mechanism ^30^. Because calcium and calcineurin signaling pathways are crucial for cell survival during cell wall stress, we examined the calcium sensitivity of the deletion strains. The *rpl40b*Δ cells were more sensitive to calcium stress, which could be a result of decreased membrane integrity. Calcium hypersensitivity was rescued by addition of calcineurin inhibitor FK506 (**Figure 4C)** ^31^. Moreover, *rpl40a*Δ *and rpl40b*Δ showed lower bulk levels of calcineurin activity compared to wild-type cells, possibly indicating an insufficient calcineurin activation (**Figure 4D**). We conclude that the loss of *RPL40B* compromises membrane integrity in general, rendering these cells sensitive to stress, such as that caused by SDS or calcium.

**Figure 4.**
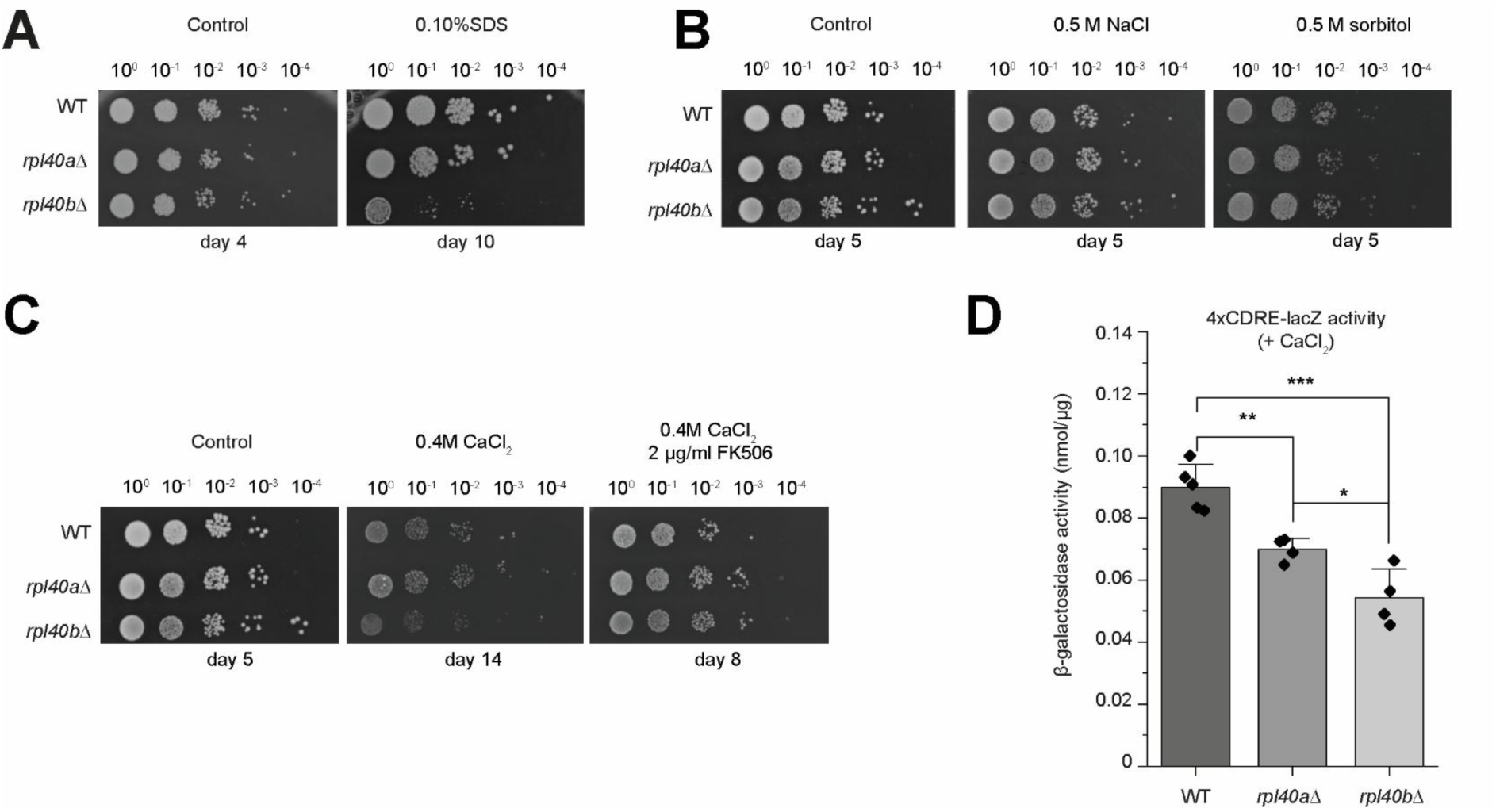
Rpl40 paralogs differentially respond to plasma membrane stress. **A.** 10-fold serial dilutions of one OD_600_ unit of yeast cells were spotted on solid plates with glycerol-containing medium without (control) or with 0.10% sodium dodecyl sulfate (SDS) added to the plate. Plates were incubated at 25°C for the indicated time. Representative images of two independent experiments are shown. **B.** 10-fold serial dilutions of one OD_600_ unit of yeast cells were spotted on solid plates with glycerol-containing medium without (control) or with 0.5 M NaCl or 0.5 M sorbitol and grown for 5 days at 25°C. **C.** 10-fold serial dilutions of one OD_600_ unit of yeast cells were spotted on solid plates with glycerol-containing medium without (control) or with 0.4 M CaCl_2_ or 0.4 M CaCl_2_ and 2 μg/ml FK506. Plates were incubated at 25°C for the indicated time. **D.** β-galactosidase activity assay of yeast strains harboring a plasmid with *lacZ* reporter gene under the control of *4xCDRE* (calcineurin-dependent response element). Strains grown in 3% glycerol + 0.05% glucose medium were incubated with 0.1 M CaCl_2_ for three hours. Data are presented as mean + SD, *n* = 2–3. Significance was determined by the two-sample *t*-test, \**p* ≤ 0.05, \*\**p* ≤ 0.01, ***p ≤ 0.001. WT, wild-type cells.

Given the link between membrane integrity, calcium homeostasis, and organelle stress signaling, we next analysed our proteomics data to determine to what extent *RPL40* paralog deletions alter the abundance of mitochondrial membrane proteins. We restricted our analysis to proteins with mitochondrial localization (**Figure 5A, B**). Pairwise comparison between *rpl40a*Δ and wild-type cells showed enrichment of proteins localizing in the inner mitochondrial membrane (**Figure 5A**).

**Figure 5.**
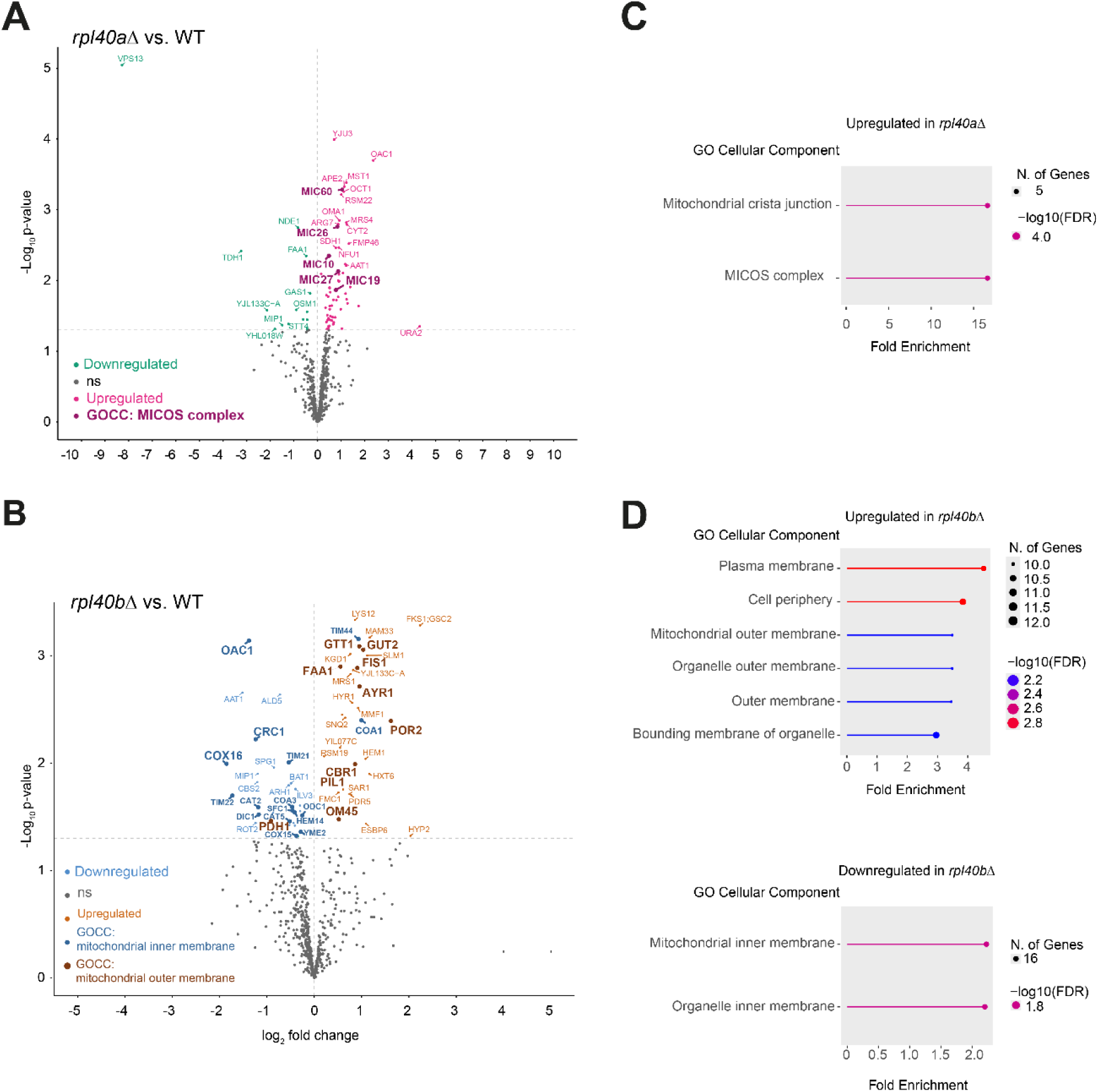
Rpl40 paralog deletions show distinct mitochondrial responses. **A, B.** Volcano plots of protein abundance changes of proteins with Gene Ontology Cellular Component (GOCC) annotation “mitochondrion” between *rpl40a*Δ vs. wild-type cells (A) and *rpl40b*Δ vs. wild-type cells (B). Significantly regulated proteins (*p* < 0.05) are highlighted in color. **C, D.** GO enrichment analysis of upregulated and downregulated genes in *rpl40a*Δ and *rpl40b*Δ strains compared to wild-type cells. Analysis using ShinyGO 0.85.1, using a customized background (1286 mitochondrial proteins) and pathway database GO Cellular Component with FDR cutoff 0.05. *n* = 4. WT, wild-type cells.

Particularly pronounced was the enrichment in components of the Mitochondrial Contact Site and Cristae Organizing System (MICOS) complex, especially Mic60, essential for the shaping of the cristae of the inner membrane and anchoring them to the mitochondrial outer membrane (MOM) by formation of contact sites with MOM proteins (**Figure 5A, C**). In contrast, deletion of *RPL40B* compared to wild-type cells showed downregulation in proteins of the inner membrane and upregulation of proteins in the outer mitochondrial membrane (**Figure 5B, D**). One example of opposite abundance of inner mitochondrial membrane protein is Oac1 (mitochondrial oxaloacetate transport protein), which increased in mitochondria of *rpl40aΔ* strain and decreased in *rpl40bΔ* strain compared to wild-type cells. Oac1 is responsible for shuttling high-energy-containing phosphoenolpyruvate (PEP) from mitochondria to the cytosol, where it contributes to the formation of triacylglycerols that are stored in lipid droplets ^32^. Indeed, a significant number of proteins downregulated in *rpl40b*Δ are members of the carrier protein family residing in the inner mitochondrial membrane (**Figure 3D**), but also Tim22, a specialized mitochondrial inner membrane protein that mediates the import and insertion of hydrophobic carrier proteins. This might indicate an adaptation of the cells to changes in the membrane composition, paralleling our findings on the sensitivity of *rpl40b*Δ cells to membrane stress.

### Rpl40 paralog-dependent regulation of mitochondrial biogenesis

Next, we aimed to understand the origin of the change in mitochondrial protein abundance dependent on the Rpl40 paralogs. First, we performed western blot analysis of mitochondrial proteins of wild-type and deletion strains from total cell extracts. We confirmed the increase of Tom20, Mia40, Pam16, and Ccp1 in *rpl40a*Δ compared to wild-type cells, whereas these proteins showed comparable levels in *rpl40b*Δ and wild-type cells (**Figure 6A**). This is in agreement with our proteomics data (**Supplementary Table 1**). To determine whether this increase is largely due to altered translation or increased transcription, we analyzed transcript levels of selected proteins. We found that *TOM20*, *MIA40*, and *CCP1* transcript levels were significantly upregulated in *rpl40a*Δ cells, consistently with the protein level (**Figure 6B**). In addition to increased transcription of mitochondrial-encoded genes, respiratory metabolism requires stabilization of mitochondrial transcripts. This process is facilitated by the mRNA-binding protein Puf3. Under fermentative conditions, Puf3 binds to mRNAs that code for mitochondrial proteins in the cytosol, facilitating decapping and deadenylation, resulting in transcript degradation and thus repression of mitochondrial biogenesis. However, when mitochondrial respiration is required, Puf3 becomes heavily phosphorylated, which leads to Puf3-dependent transcript stabilization and translocation to the mitochondrial outer membrane. To determine whether altered Puf3 regulation contributes to the mitochondrial biogenesis phenotype observed in the *RPL40* deletion strains, we generated HA-tagged Puf3 strains and analyzed the phosphorylation status of Puf3 using SDS-PAGE/Western blotting electrophoresis with Phos-tag, to detect phosphorylated proteins, which show a distinct mobility shift due to their interaction with Phos-tag (**Figure 6C**). Under glycerol growth conditions, both the *rpl40a*Δ and *rpl40b*Δ strains exhibited an increased abundance of slower-migrating Puf3 fractions compared to wild-type cells, which is consistent with elevated Puf3 phosphorylation in mutant strains; however, the overall pattern of protein bands is substantially distinct, showing that *rpl40b*Δ resembles the wild-type, while *rpl40a*Δ shows higher content of slower-migrating protein bands, that might suggest that in this particular strain Puf3 is hyper-phosphorylated. Total protein levels of Puf3 remained comparable between strains (**Figure 6D**). To analyse potential alterations in Puf3-dependent mediation of mitochondrial transcript levels in the *RPL40* deletion cells, we performed a medium-switch experiment, in which cells were initially grown on glucose-containing medium to logarithmic growth phase, washed and cultured in glycerol-containing medium for 1.5, 3 or 20 hours (**Figure 6E-I**). *COX17* mRNA is a target of Puf3 regulated by a carbon-specific mechanism ^33^. In dextrose/glucose conditions, *COX17* mRNA decay is regulated by Puf3. Even though Cox17, is not classified as a cis target protein by Lapointe et al.^33^, we see low Cox17 protein levels in wild-type cells in glucose as a carbon source. However, in the absence of *PUF3*, the protein is strongly produced in glucose conditions and fails to be induced when cells are shifted to glycerol. A similar pattern is observed for Cox17 in the *RPL40* deletion strains, with a more pronounced increase in Cox17 expression in *rpl40a*Δ compared to both wild-type and *rpl40b*Δ cells (**Figure 6E, F**). Ccp1 is a trans-target of Puf3, whose protein abundance is indirectly dependent on Puf3p and Ccp1 protein abundance is generally decreased in *puf3*Δ. However, it is strongly increased in *rpl40a*Δ compared to wild-type and *rpl40b*Δ cells, already under glucose conditions (**Figure 6E, G**). Similarly, *MIA40* mRNA is Puf3-bound mRNA, and Mia40 protein is highly increased in *rpl40a*Δ strain under glucose conditions (**Figure 6H, I**). Thus, although Puf3 phosphorylation is elevated in *rpl40a*Δ and *rpl40b*Δ strains, increased mitochondrial protein abundance is only observed in the *rpl40a*Δ strain, probably due to higher phosphorylation level of Puf3. This suggests that the loss of Rpl40a in particular results in the uncoupling of transcriptional and translational control of mitochondrial biogenesis and appears to be independent of the metabolic state of the cells.

**Figure 6.**
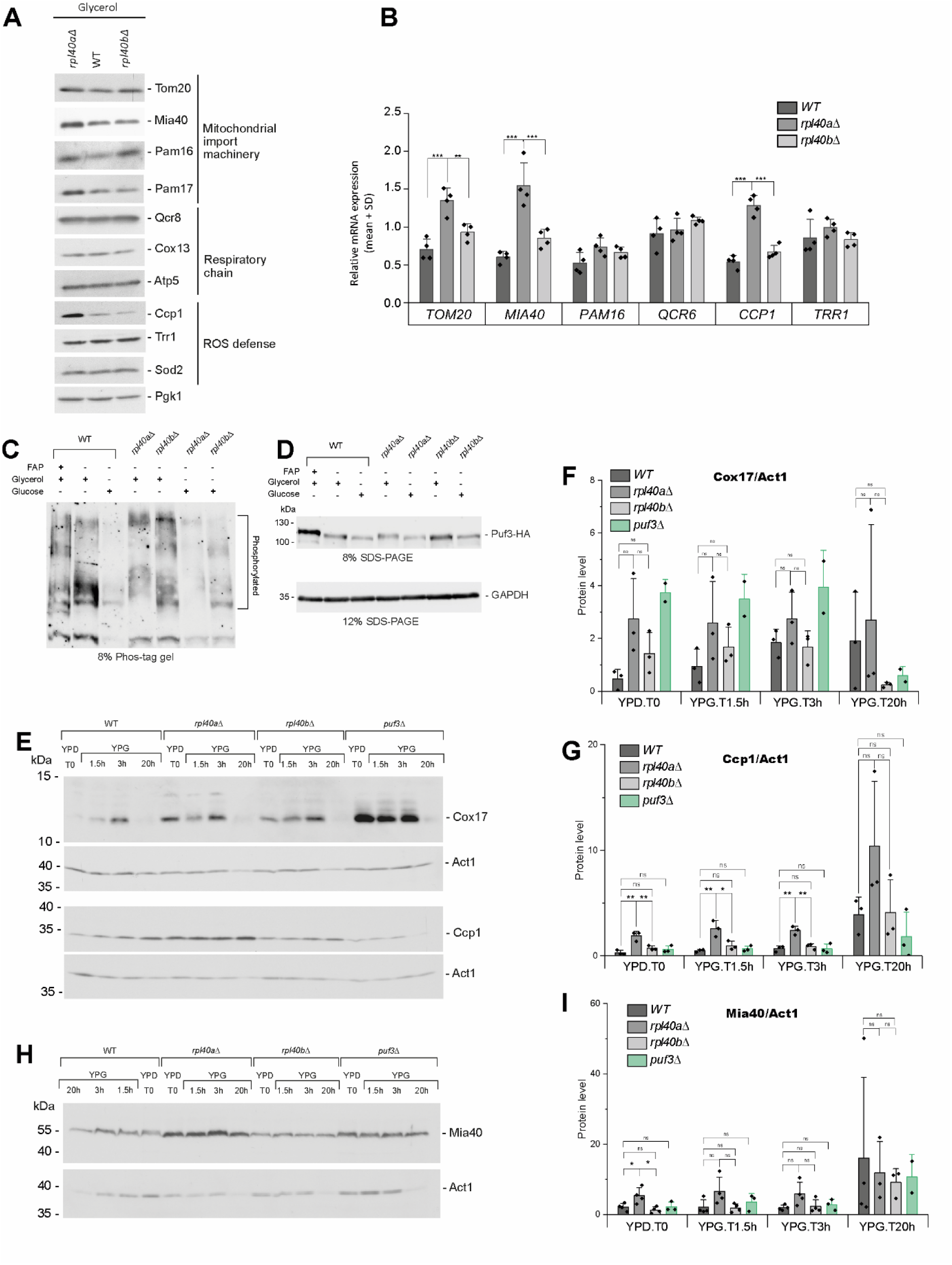
Rpl40 paralogs differentially control mitochondrial biogenesis through transcriptional and Puf3-dependent mechanisms. **A.** Western blot analysis of *RPL40* deletions and wild-type (WT) strain. Cells were grown at 28°C on glycerol-containing medium. Total protein extracts were separated on SDS-PAGE and analysed by western blot using specific antibodies. **B.** Transcript levels of wild-type (WT), *rpl40a*Δ, and *rpl40b*Δ strains grown at 28°C on glycerol-containing medium. Transcript levels of indicated genes were calculated based on the concentration of amplified products and relative to the geometric mean of the *ACT1*, *ALG9* and *TDH1*. Graph represent the mean + SD, *n* = 4. \*\**p* ≤ 0.01, \*\*\**p* ≤ 0.001. **C.** Analysis of the phosphorylation state of Puf3 using SDS-PAGE/Phos-tag coupled with western blot; the electrophoretic mobility of phosphorylated proteins was evaluated upon addition of phos-tag. Yeast cells were grown on glucose or glycerol-containing medium, cells were harvested at the logarithmic growth phase; 20 μg of total cell extract was loaded per sample and detection was performed based on anti-HA antibodies; FAP – treatment of the cell extract with alkaline phosphatase. **D.** Total cells extract was separated by SDS-PAGE and analysed by western blot with anti-HA to detect Puf3-HA and anti-GAPDH, which serves as loading control. **E, H**. Western blot assay assessing Cox17, Ccp1 and Mia40 proteins levels in wild-type (WT), *rpl40a*Δ, and *rpl40b*Δ cells. Yeast cells were grown on glucose-containing medium (YPD) to the logarithmic growth phase, washed and resuspended in glycerol-containing medium (YPG). Cells were harvested at the indicated time points. Total protein extracts were separated by SDS-PAGE and analysed by western blot using specific antibodies. Representative images of 3–4 independent replicates. **F, G, I**. Quantification of western blot signals of chosen proteins relative to Act1 control protein. Data are presented as mean + SD, *n* = 2–4. WT, wild-type cells; T0 is time just before the shift, T1.5h is 1.5 hours after the shift, T3h is 3 hours after the shift and T20h is 20 hours after the shift.

### Reduced fatty acid content upon *RPL40* paralog deletions does not alter mitochondrial fatty acid metabolism

Given the changes in abundance of proteins residing in the mitochondrial membranes, we analysed the fatty acid metabolism to detect possible upstream causes linked to the observed alterations in mitochondrial membrane proteins. The content of 14 fatty acids was analysed from mitochondria and culture medium (**Supplementary Figure 4A, Supplementary Table 2**). Culture medium of *rpl40b*Δ contained increased concentration of capric acid (C10:0) and decreased concentration of stearic (C18:0) and lignoceric (C24:0) acids compared to *rpl40a*Δ. However, in the case of mitochondria, the total level of unsaturated fatty acids (UFA) was increased in *rpl40a*Δ, especially significantly C18:2 and C22:1, compared to wild-type cells. We found that both deletion strains had significantly lower levels of medium-chain saturated fatty acids [octanoic acid (C8:0)] and long-chain saturated fatty acids [myristic acid (C14:0), arachidic acid (C20:0)] and one very long-chain fatty acid, lignoceric acid (C24:0).

In yeast, C14:0 and C24:0 are components of phospholipids essential for the physical properties of biological membranes, including those in mitochondria. While C24:0 serves a vital structural and organizational role within mitochondria and cell membranes, C14:0 is rather a minor component of yeast membranes. Both long-chain saturated fatty acids have a structural role rather than being used as an energy source. In contrast, C8:0 synthesized within mitochondria serves as a crucial precursor for the production of lipoic acid, an essential cofactor for vital enzymes in the Krebs cycle and respiratory chain. To elucidate whether the altered mitochondrial function of the deletion mutants depends on the modification of key proteins modified with lipoic acid, we analysed expression levels of key enzymes of the mitochondrial fatty acid synthesis pathway (**Supplementary Figure 4B**). We found that in *rpl40a*Δ the expression of the mitochondrial acyl carrier protein *ACP1* was increased, and this correlated with increased Acp1 protein level in mitochondria (**Supplementary Table 1**). Acp1 protein localizes to the mitochondrial matrix and is essential for biosynthesis of octanoate (conjugate base of caprylic acid), which is a precursor to lipoic acid, indicating a potential compensation mechanism for low levels of the fatty acid C8:0 in the deletion mutants (**Supplementary Figure 4C**). Moreover, Acp1 is also a cis Puf3 target protein. Changes in the expression of other enzymes in the pathway were not observed apart from a minor increase in *LIP2* in *rpl40b*Δ. Finally, we analysed the expression of *GCV3* (glycine cleavage system H protein, mitochondrial), *LAT1* (dihydrolipoamide acetyltransferase component of pyruvate dehydrogenase complex), and *KGD2* (alpha-ketoglutarate dehydrogenase subunit E2), proteins directly or indirectly involved in the Krebs cycle, and their lipoylation state. Apart from a minor increase in the transcript levels of *KGD2*, we did not observe any changes in the protein levels based on our proteomics analysis (**Supplementary Figure 4C, Supplementary Table 1**). Moreover, levels of the lipoylated form of the proteins were comparable to the levels in wild-type cells (**Supplementary Figure 4D**). Thus, we conclude that low levels of cellular ATP in the *RPL40* paralog deletions are not caused by significant changes in the mechanisms that supply the electron transport chain with chemical energy in the form of NADH and FADH2 (**see Figure 2F**). This analysis also supports our data on the functionality of the respiratory chain in *RPL40* deletion strains (**see Figure 2E**). Moreover, Acp1 regulates the assembly of the electron transport chain (ETC) complexes ^34^. Nevertheless, mitochondrial fatty acid synthesis can be involved in the production of other fatty acids and not only octanoic acid, so alterations in enzymes of this pathway can lead to imbalance levels of fatty acids by adversely affecting the pool of lipids in the cell with broad consequences for membrane integrity.

### *RPL40* paralogs deletions show pronounced differences in triglyceride content

To evaluate differences in membrane composition, we performed lipidomic profiling on total cell extracts and isolated mitochondria from wild-type and *RPL40* deletion strains, each in four biological replicates (**Supplementary Table 3, Figure 7, Supplementary Figures 5 and 6**). In total cell extracts, we detected 817 lipids across 26 lipid classes, while mitochondrial samples contained 837 lipids belonging to 23 classes. Principal component analysis (PCA) from total cell extract samples revealed clear separation between the wild-type and the deletion strains, while also showing that the two deletion strains form distinct clusters from each other (**Supplementary Figure 5A**). This contrasts with mitochondrial lipidomes, where the *rpl40b*Δ strain clustered closely with the wild-type, whereas the *rpl40a*Δ strain formed a clearly distinct group (**Supplementary Figure 6A**). Most lipid classes were altered in total lipidomes of *RPL40* deletion strains **(Supplementary Figure 5B).** However, lipid classes in isolated mitochondria showed more pronounced changes *rpl40aΔ* cells compared to the wild-type (**Supplementary Figure 6B**). *RPL40A* deletion strain displayed a marked accumulation of triacylglycerols (TAG) in total lipidome together with elevated cardiolipin levels in mitochondria (**Figure 7A**), whereas the *RPL40B* deletion strain showed a pronounced reduction in triacylglycerols without detectable changes in cardiolipin abundance compared with wild-type strain (**Figure 7B**). Cardiolipin is an important cone-shaped non-bilayer phospholipid of the inner mitochondrial membrane that induces the formation of high-curvature regions essential for cristae organization, and its synthesis depends on the cardiolipin synthase, Crd1. Our proteomics analysis revealed that Crd1 protein level in *rpl40a*Δ cells is significantly increased compared to *rpl40b*Δ strain (log2 FC = 0.7, **Supplementary Table 1**), which could support higher cardiolipin synthesis in *rpl40a*Δ. Analysis of phospholipid classes, including phosphatidic acid (PA), phosphatidylinositol (PI), phosphatidylserine (PS), phosphatidylethanolamine (PE) and phosphatidylcholine (PC), according to the saturation (the number of double bonds) and total number of carbons in the acyl-tails revealed distinct remodeling patterns. *RPL40A* deletion strain exhibited reduced biosynthesis of PI and PC molecular species containing long, unsaturated fatty acids (**Figure 7C**). On the other hand, *rpl40b*Δ strain showed decreased production of PI and PC phospholipids enriched in long, saturated fatty acids (**Figure 7D**). Together, these results indicate that the *RPL40* paralog deletions differentially affect neutral lipid storage and mitochondrial membrane lipid composition, as well as the balance of acyl-chain saturation and length in phospholipids. In particular, our data indicate that PC, the most abundant phospholipid in membranes ^35^, contains reduced levels of long saturated fatty acids in the *rpl40b*Δ strain, which can change the physical properties of membranes, including thickness, fluidity, or membrane curvature, as well as the function of membrane proteins, giving a reasonable explanation for the cells’ hypersensitivity to membrane stress.

**Figure 7.**
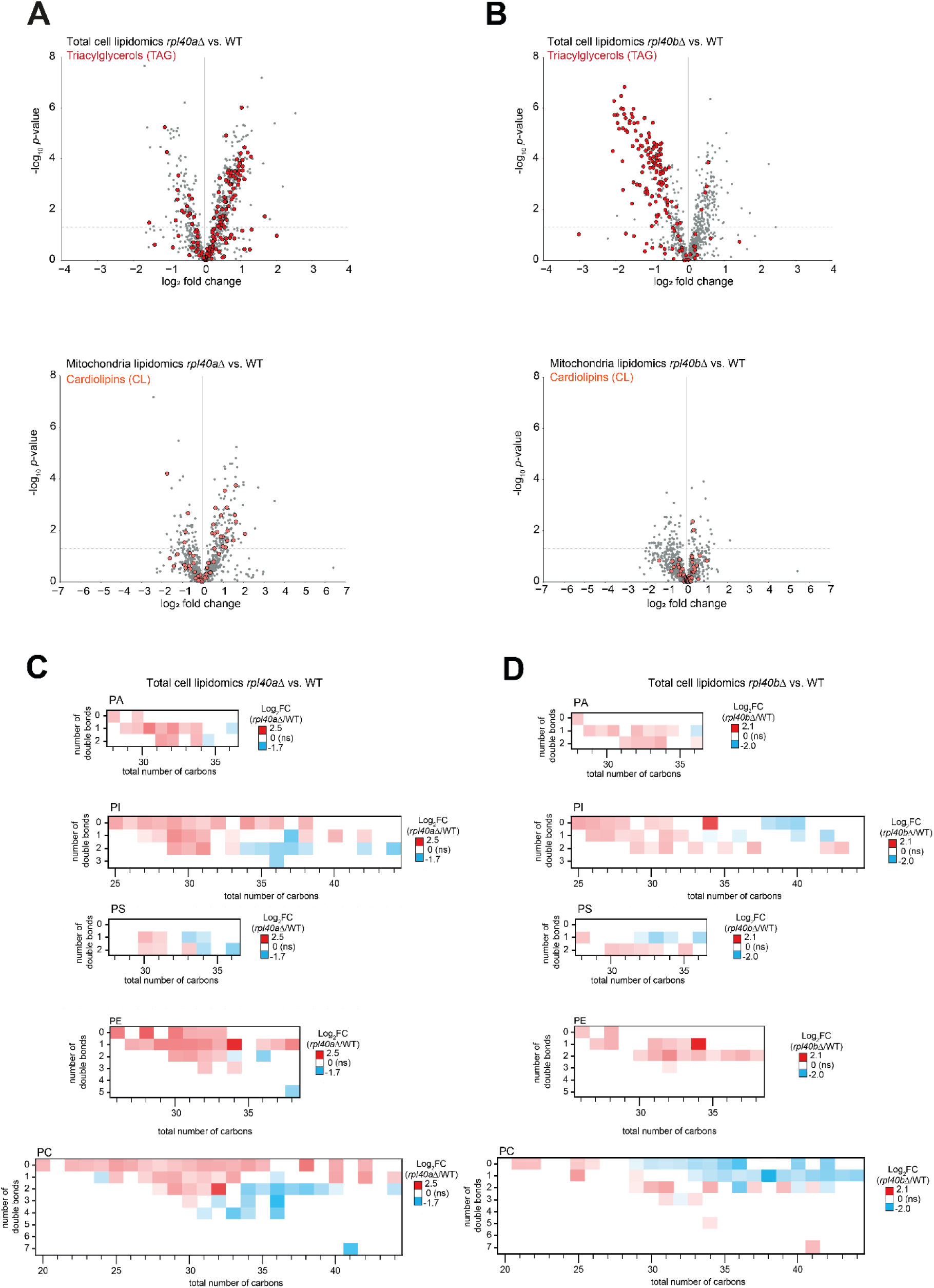
Rpl40 paralogs differentially impact the cellular lipidome. **A, B.** Volcano plots of triacylglycerols (TAG) alterations in total cell lipidome and cardiolipins (CL) alterations in mitochondrial lipidome of *rpl40a*Δ strain (A) and *rpl40b*Δ (B) compared to the wild-type cells (WT), *n* = 4. A dashed horizontal line indicates *p* value 0.05. **C, D.** Heatmaps of log_2_ fold change (FC) [(C) *rpl40a*Δ vs. WT; (D) *rpl40b*Δ vs. WT] of significantly changed phospholipid classes: phosphatidic acid (PA), phosphatidylinositol (PI), phosphatidylserine (PS), phosphatidylethanolamine (PE), and phosphatidylcholine (PC), according to their molecular species. Saturation and length of PA, PI, PS, PE, and PC species are given by the number of double bonds or the total number of carbons in the acyl tails, respectively. If the same species had different retention times, one of them was selected, provided that their fold changes were similar. If their fold changes were opposite, then that species was not included in the heatmap.

## DISCUSSION

Mitochondrial biogenesis is largely dependent on nuclear-encoded genes and therefore is driven by efficient synthesis of mitochondrial proteins by cytosolic ribosomes followed by their import into the organelle. Growing evidence indicates that ribosomal paralogs and variable ribosomal protein composition contribute to the functional heterogeneity of the cellular translation machinery. Whether specialized cytosolic ribosomes act as upstream regulators of mitochondrial function by controlling the translation of nuclear-encoded mitochondrial proteins remains an important question with relevance for human ribosomopathies, which are often accompanied by mitochondrial dysfunction ^36^. In this study, we found that Rpl40/eL40 paralogs play differential roles in the regulation of mitochondrial function and cellular homeostasis in the budding yeast *S. cerevisiae*. Rpl40b promotes the exaggeration of mitochondrial biogenesis and increases lipid storage and membrane integrity. By contrast, a decrease of cytosolic translation under respiratory growth conditions is Rpl40a dependent, with concomitant decrease in lipid storage and membrane integrity but increase of mitochondrial mRNA translation. Here, our data diverge from previous published work. While our data suggest that the complementary functions of the Rpl40 paralogs support mitochondrial morphology and biogenesis, previous findings in yeast deletion strains lacking specific RP paralogs either support or do not support mitochondrial function. In this respect, Rpl1b, Rpl2b, Rps11a and Rps26b were required for normal mitochondrial function and morphology, whereas deletion of the corresponding paralog did not result in respiratory deficiencies ^11^. This opens the possibility that Rpl40 paralogs can support adjustment of mitochondrial function to cellular metabolic demands.

Our data are limited in a way that they do not provide direct proof of the potential selective translation of mRNAs depending on the Rpl40 paralogs. However, evidence in mammalian cells suggests a role in selective translation depending on eL40 (encoded by *UBA52*). Here, bulk cellular translation of rapidly proliferating cells does not depend on rpL40^21^, which is consistent with our analysis of newly synthesized proteins under fermentative conditions. Depletion of rpL40, however, affected a set of selected mRNAs, including those involved in stress responses. In this context, the opposing phenotypes observed in the *rpl40a*Δ and *rpl40b*Δ strains in this study may reflect differential translation of subsets of mRNAs involved in mitochondrial biogenesis, membrane remodeling, or stress signaling. A single gene encodes rpL40 in mammalian cells, and no isoforms have been reported to this day, raising the question of how selective translation can be achieved. Recent work demonstrated that specificity might be achieved in mammalian cells not through paralog specialization but through dynamic remodeling of ribosomal composition by recruitment of additional rpL40 protein to specific ribosomal populations. During viral entry, rpL40 is recruited to a noncanonical position on the small subunit of 80S ribosomes near the mRNA entry channel, generating specialized ribosomes that preferentially associate with viral mRNAs and promote efficient viral protein synthesis ^22^. Notably, the increased rpL40 occupancy on ribosomes was triggered by membrane fusion between the virus and host. Moreover, serum starvation of cells also resulted in increased occupancy of ribosomes with rpL40. Loss of rpL40 during serum starvation resulted in downregulation of mRNAs of essential metabolic and intracellular transport pathways. These included mRNAs involved in autophagosome-lysosome fusion and endosome recycling, such as components of Endosomal Sorting Complex Required for Transport (ESCRT) machinery ^22^. Membrane repair pathways involving the ESCRT machinery are also important for both viral budding and plasma membrane damage responses, raising the possibility that rpL40-dependent translational programs may influence conserved membrane remodeling pathways across eukaryotes. Cellular or environmental conditions that would trigger the selective incorporation of either Rpl40 paralog into the yeast ribosome remain to be shown. To this end, our data support a distinct but complementary function of both Rpl40 paralogs during respiratory growth conditions since deletion of either paralog showed slow growth and defects in mitochondrial morphology. However, nutrient depletion, transitioning to the stationary growth phase, or mitochondrial dysfunction could trigger the need for adjusting translational output ^37–40^. Stress conditions commonly induce global repression of cytosolic translation while selectively maintaining or transiently enhancing translation of mitochondrial encoded genes ^41,42^, a phenotype that is manifested in the *rpl40b*Δ. Mitochondrial function is closely linked with lipid metabolism since mitochondrial biogenesis, oxidative phosphorylation, and stress resistance require extensive membrane and lipid remodeling ^43–45^. The pronounced reduction in triacylglycerol levels in the *rpl40b*Δ strain may reflect impaired lipid synthesis and storage, but it may also indicate increased mobilization of storage lipids to sustain membrane remodeling under chronic stress. Recent work demonstrated that mitochondrial stress induces active mobilization of triacylglycerol stores, with liberated fatty acids being redirected toward cardiolipin synthesis and mitochondrial membrane biogenesis during stress recovery ^46^. Thus, depletion of TAGs in *rpl40b*Δ cells may reflect sustained lipid consumption required to support chronic mitochondrial remodeling and respiratory adaptation. In contrast to the TAG depletion observed in *rpl40b*Δ cells, the increased TAG accumulation detected in *rpl40a*Δ cells may represent a protective adaptation to mitochondrial dysfunction rather than enhanced lipid storage capacity itself. Numerous studies have shown that mitochondrial stress promotes lipid droplet biogenesis and TAG accumulation as a mechanism to buffer excess fatty acids, reduce lipotoxicity, and protect cellular membranes from oxidative damage ^47,48^. In yeast, the transition from fermentative to respiratory metabolism is also accompanied by a rapid induction of lipid droplet biogenesis ^49–51^. These findings are consistent with the phenotype of *rpl40a*Δ cells, in which excessive production of nuclear-encoded mitochondrial proteins triggers translational imbalance leading to mitochondrial stress and ROS accumulation, which in turn promotes lipid remodeling and diversion of fatty acids into TAG storage.

Moreover, changes in lipid composition of mitochondrial membranes, including the abundance of glycerophospholipids, unsaturated fatty acids, as well as a functional mitochondrial fatty acid synthesis pathway, can affect mitochondrial DNA stability and increase the frequency of petite cells ^52^. Cardiolipins are important for stabilization of mtDNA ^53^, thus, the increased levels of cardiolipin species in *rpl40a*Δ cells may prevent mtDNA depletion or mutation and decrease the formation of respiration-deficient petite cells. However, mtDNA loss is also a mechanism of adaptation during oxidative stress ^54^. Thus, reduced ability to form respiratory-deficient cells may explain the decreased viability of *rpl40a*Δ cells during oxidative and acetic acid stress.

In summary, our data identified the Rpl40 paralogs as potential upstream regulators of mitochondrial biogenesis and lipid homeostasis. Given the growing evidence that ribosome heterogeneity contributes to selective translational control, the distinct mitochondrial and lipid phenotypes observed in the *rpl40a*Δ and *rpl40b*Δ strains support a model in which Rpl40-dependent ribosomal populations coordinate the synthesis of proteins required for respiratory adaptation, membrane remodeling, and storage lipid metabolism during cellular stress responses.

## MATERIALS AND METHODS

### Yeast strains and growth conditions

Budding yeast, *Saccharomyces cerevisiae* strains derived from BY4741 background were used in this study and are listed in **Supplementary Table 4**. All yeast strains were maintained on YPD solid plates (1% yeast extract, 2% peptone, 2% glucose, and 2% agar). Depending on the experimental setup, yeast cells were grown in the indicated medium and temperature to the logarithmic growth phase (optical density OD600 ∼0.6–1), unless stated otherwise.

To analyse growth of strains in liquid medium, cells were cultured in YPD (1% yeast extract, 2% peptone, 2% glucose), then diluted to OD600 ∼0.2 in YPD or YPG (1% yeast extract, 2% peptone, 3% glycerol, pH 5.2) or cultured in YPD until the logarithmic growth phase, and then washed with sterile water and transferred to YPG medium. Next, 200 μl of cell suspension was transferred to a 96-well plate and incubated at 28°C for 48 –72 hours with continuous orbital shaking. The absorbance at 600 nm was measured every 60 minutes using Agilent BioTek Synergy H1 plate reader (Agilent Technologies, Inc). To analyse the growth of strains in solid medium, cells were diluted ten-fold in sterile water from OD600=1 (dilution 0). Two-microliter drops of each dilution were pipetted on solid medium plates with indicated carbon sources and incubated at specified temperatures for indicated time.

### Generation of HA-tagged PUF3

The 3xHA-tag insert together with hygromycin resistant cassette was amplified by PCR from the empty vector pYM24 (Cat. no P30236, Euroscarf). Primers used for amplification were designed to add flanks at the N- and C-terminus of the PCR product homologous to the C-terminus of the *PUF3* gene and to the downstream region of the gene, respectively (**Supplementary Table 5**). The purified PCR product was transformed into BY4741 (WT) yeast strain and selection of transformants was performed on YPD agar plates supplemented with hygromycin at the concertation of 300 μg/ml. The presence of C-terminal 3xHA-tag at *PUF3* gene was confirmed by PCR and by Sanger sequencing. The resulting strain yMS032 was used as background to delete *RPL40A* and *RPL40B*. Deletion was achieved trough homologous replacing the genes with URA3 selection marker, which was amplified from the empty vector pRS306. Primers used for amplification of deletion cassette were designed to add flanks at the N- and C-terminus of the PCR product homologous to the up- and downstream region of the gene (**Supplementary Table 5**). Deletion of each gene was confirmed by PCR and Sanger sequencing. All resulting strains are listed in Supplementary Table 4.

### Petite frequency assay

The frequency of forming petite colonies was determined according to the modified protocol of ^55,56^. Yeast strains were grown overnight in liquid YPG medium shaking 150 rpm, then diluted to an OD600 of ∼0.02 and cultured in YPG liquid medium for 48 hours at 28°C with orbital shaking 150 rpm. Next, ten-fold serial dilutions of cells from starting OD600=1 were spotted on YPD solid plates and incubated for two days at 25°C. A single colony from each plate was picked and resuspended in 20 ml of sterile water. Then, 500 μl of the solution was diluted additionally with 600 μl of sterile water and 10 μl of the suspension were plated evenly onto a YPG plate containing 0.05% glucose. Plates were incubated for seven days at 25°C and scanned. Next, 15 ml of solution containing 0.6% agar, 0.5% glucose, 9.8 mM Tris-HCl, pH 8.0, and 0.36% 2,3,5-Triphenyltetrazolium chloride (cat. no T8877-25G, Sigma Aldrich) was poured evenly over the plates. After 25 minutes of incubation at room temperature, the plates were scanned again. The number of total colonies before tetrazolium overlay, and the number of white and red colonies after tetrazolium overlay, were counted using open-source software DotDotGoose ver.1.7.0 (biodiversityinformatics.amnh.org/open_source/dotdotgoose/). Petite frequency was calculated as a percentage of the number of white (petite) colonies to the total number of colonies on a plate.

### Translation assays

#### Cytoplasmic translation

Yeast cells were grown at 28°C to logarithmic growth phase (OD600 ∼0.6–1) on minimal synthetic medium [0.67 g/l yeast nitrogen base, 0.79 g/l complete supplement amino acid mixture (CSM)] containing different carbon sources: 2% (v/v) glucose or 3% (v/v) glycerol. Cells were starved in methionine-free minimal medium with the corresponding carbon source for 15 minutes. The proteins were radiolabeled using [^35^S]-labeled methionine (cat. no. SRM-01H, Hartmann Analytic, Braunschweig, Germany) for 30 minutes at a final concentration of 6 μCi ml^−1^. After harvesting, yeast cells were washed once with double-distilled H2O (ddH2O). Total protein extracts were prepared based on alkaline lysis for 10 minutes on ice, and proteins were precipitated with a final concentration of 10% trichloroacetic acid (TCA). The extracted proteins were resuspended in Laemmli buffer containing 50 mM 1,4-dithiothreitol (DTT, cat. no 6908.2, Carl Roth GmbH + Co.). Proteins were denatured for 15 minutes at 65°C, analysed by SDS-PAGE and subsequent autoradiography using Fujifilm FLA-7000 (GE Healthcare, Bio-Sciences AB, Uppsala, Sweden).

#### Mitochondrial translation

Yeast cells were grown in YPG medium at 24°C to the logarithmic growth phase. Ten OD600 units of yeast cells were harvested by centrifugation (5 minutes, 3000xg, RT). The pellet was resuspended in methionine-free medium supplemented with 3% glycerol, and the yeasts were cultured for one hour at 24°C. Next, 6 OD600 units of yeast cells were spun down, and the pellet was resuspended in 500 µl of minimal synthetic medium [0.67 g/l yeast nitrogen base, 0.77 g/l CSM lacking methionine] containing 3% glycerol. To block cytoplasmic protein synthesis, cycloheximide (CHX) was added to a 0.225 mg/ml final concentration. Cells were incubated for 5 minutes at 24°C, and [^35^S]-labelled methionine (cat. no. SRM-01H, Hartmann Analytic, Braunschweig, Germany) was added for 30 minutes at a final concentration of 50 μCi ml^−1^. Next, yeast cells were harvested by centrifugation (1 minute, 10,000xg, RT). The pellets were resuspended in 75 µl of lysis buffer (1.85 M NaOH, 14% (v/v) beta-mercaptoethanol, 10 mM PMSF). Proteins were precipitated by the addition of 1 ml of 25% TCA. After centrifugation (5 minutes, 20,000xg, RT), supernatants were discarded and the pellets were washed with 2 M Tris base. Samples were spun down (1 minute, 20,000xg, RT), and the pellets were resuspended in 50 µl of Laemmli buffer containing 50 mM DTT. Proteins were separated using SDS-PAGE in running buffer (50 mM Tris, 384 mM glycine, 0.1% SDS) adjusted with HCl to pH 8.3. The gel was blotted on PVDF membrane, and dried membrane was exposed in Storage Phosphor Screen (GE Healthcare) for 7 days before signal detection using phosphoimager Fujifilm FLA-7000 (GE Healthcare, Bio-Sciences AB, Uppsala, Sweden).

#### Isolation of RNA from yeast cells, cDNA synthesis, and quantitative PCR

Twenty OD600 units of exponentially grown cells were collected for isolation of total RNA and cDNA synthesis, as explicated previously ^57^. Primers were designed using SGD’s Primer Design tool and Primer-BLAST ^58^ and are presented in **Supplementary Table 5**. Each reaction was performed in technical duplicate/triplicate by using RT PCR Mix SYBR C (cat. no 2008-1000, A&A Biotechnology) on Roche LightCycler 480 System. The reaction program included an initial denaturation at 95°C for 5 minutes, and 40 cycles consisting of following steps: 95°C for 30 s, 58°C for 20 s, and 72°C for 20 s. To ensure the specificity of amplification, melting curve analysis was performed following each experiment. For each set of primers, negative control and cDNA dilutions for the standard curve were loaded on a plate. The amplification efficiency of each primer pair was assessed, and the relative expression of the gene was calculated using the adjusted 2^-ΔΔCT^ method ^59^. Relative expression of each gene to the geometrical mean of reference genes (*ACT1, ALG9*, and *TDH1*) was calculated for at least three independent biological replicates. Results were normalized to the WT group.

#### Mitochondrial morphology assessment

Mitochondrial morphology was assessed using the expression of mitochondria-targeted green fluorescent protein (GFP) under a fluorescence microscope. Yeast were transformed with the pYX122-mtGFP plasmid (Addgene plasmid #45048, a gift from Benedikt Westermann) and grown in histidine-free minimal medium supplemented with 2% glucose or 3% glycerol to logarithmic growth phase (OD600 0.6–1). One OD600 unit was spun down, dissolved in 100 μl PBS, and placed on a coverslip. Fluorescence images of yeast cells were obtained using Carl Zeiss Axio Imager M2 microscope. Green fluorescence from 488 nm was measured using a filter under 100x magnification. The Z-stack image series consisted of 0.3-0.588 µm slice intervals were taken in two biological replicates. Obtained images were processed using Fiji ImageJ with MINA plug-in for mitochondria morphology analysis, as described previously ^60^. Non-parametric statistical test (Kruskal-Wallis) was applied to assess differences in the mean number of mitochondria network branches.

#### Transmission electron microscopy

Yeast cells cultured in glycerol containing medium until OD600 0.6–0.7 were cryo-immobilized without cryoprotectant by high-pressure freezing using a Leica EM ICE high-pressure freezer. Samples were loaded into aluminum carriers (100 µm cavity) and immediately frozen. Freeze substitution was performed in 1% osmium tetroxide (Agar Scientific) and 0.1% uranyl acetate (CHMES) in acetone using a Leica AFS freeze substitution unit. Samples were maintained at −90°C for 12 hours, followed by −84°C for 72 hours, and then gradually warmed to −30°C for 1 hour, −15°C for 1 hour, and finally 0°C. After freeze substitution, samples were washed three times in anhydrous acetone and infiltrated with Embed-it™ Low Viscosity Epoxy Resin (Polysciences) using a graded resin-acetone series (1:3, 1:1, 3:1, and pure resin). Samples were incubated in resin overnight and embedded in fresh resin. Polymerization was carried out at 60°C for 48 hours. Ultrathin sections (70 nm) were cut using a Leica EM UC7 ultramicrotome, collected on 200-mesh nickel grids (Agar Scientific), and post-stained with 1% lead citrate (Micro to Nano). Sections were examined using a Tecnai T12 BioTwin transmission electron microscope (FEI, Hillsboro, OR, USA) equipped with a 16-megapixel TemCam-F416 camera (TVIPS GmbH) at the Microscopy and Cytometry Core Facility, IN-MOL-CELL Infrastructure, International Institute of Molecular and Cell Biology in Warsaw (IIMCB), Warsaw, Poland.

#### Measurement of total ATP levels

Twenty OD600 units of yeast cells were harvested and washed once in lysis buffer (20 mM Tris acetate, pH 7.75, 2 mM EDTA). The yeast cell pellet was resuspended in lysis buffer and incubated at 90°C for 5 minutes with agitation. The lysate was cleared by short centrifugation, and the cleared lysate was analysed according to the manual instruction of ENLITEN ATP Assay kit (cat. no FF2000, Promega).

#### Measurement of reactive oxygen species

For stationary phase measurement BY4741, *rpl40a*Δ, *rpl40b*Δ strains were grown from OD600=0.2 in minimal CSM medium with 2% glucose for 48 hours, at which time the glucose levels were depleted by dividing cells. Cells were cultured at the stationary phase at OD600 7–9. One OD600 unit of yeast cells was collected and washed with PBS (137 mM NaCl, 12 mM phosphate, 2.7 mM KCl, pH 7.4). Yeast cells were resuspended in PBS containing 5 µM dihydroethidium (cat no. D7008, Sigma-Aldrich). The following controls were prepared for each condition: PBS only, yeast cells resuspended in PBS without dye, and PBS only with dye. Samples were incubated for 40 minutes at room temperature in the dark before measurement. Fluorescence signal was measured at an excitation wavelength of 535 nm and an emission wavelength of 635 nm using a plate fluorometer (SpectraMax iD3, Molecular Devices). Fluorescence signals were collected at 5-minute intervals for 2 hours. Fluorescence units from each sample were corrected for autofluorescence and autoxidation of the dye in PBS. Corrected data are presented as mean + SD of the slope value calculated with the SLOPE() formula in MS Office Excel.

#### Calcineurin activity measurements

BY4741 (WT), *rpl40aΔ, rpl40bΔ* strains were transformed with plasmid PAMS366 containing four copies of calcineurin-dependent response element (4xCDRE) placed upstream of a *lacZ* reporter gene. Cells were grown in synthetic medium lacking uracil [0.77g/l complete supplement mixture of amino acids without uracil (CSM-URA), 6.7 g/l yeast nitrogen base (YNB), 3% glycerol, and 0.05% glucose] until the logarithmic phase and then treated with 0.1 M CaCl2 or sterile water for 3 hours at 28°C. 25 OD600 units of cells were collected, washed with sterile water, and stored at −80°C. The β-galactosidase activity was measured as described previously ^61^. Briefly, cells were resuspended in 50 mM phosphate buffer (KPi, a mixture of monobasic dihydrogen phosphate and dibasic monohydrogen phosphate, pH 7.0) and frozen with glass beads (cat. no G9268-250G, Sigma-Aldrich) using liquid nitrogen. After thawing at room temperature, lysates were vortexed for 6 minutes at 4°C, and centrifuged (15,000xg, 4 minutes, 4°C). Supernatants were transferred to new tubes for further analysis. Protein concentration of cell extracts was measured by the Bradford method using ROTI®Quant (cat. no K015.3, Carl Roth) and Bovine Serum Albumin (BSA) as calibrating protein (Thermo Scientific™, cat. no 23209). Cell extracts (5-25 μl) were mixed with 1 ml buffer (60 mM Na2HPO4, 40 mM NaH2PO4, 10 mM KCl, 1 mM MgSO4). The 200 μl of substrate 0.4% o-nitrophenyl-β-D-galactopyranoside (ONPG, BioShop Canada Inc., cat. no ONP301.5) in 50 mM Tris-HCl, pH 8.0, was added to start the reaction. When the solution became yellow, the reaction was quenched by adding 500 μl 1M Na2CO3. The amount of the product was measured at an absorbance of 420 nm. The β-galactosidase activity in one minute of the reaction was normalized to 1 μg of protein in the cell extract. Two biological repetitions of each two different transformants per strain were measured in technical duplicates.

#### Quantification of mitochondrial DNA

Genomic DNA isolation was performed according to Tahmaz et al. ^62^. Briefly, cells were grown in YPG medium until the logarithmic growth phase. 20 OD600 units were collected, the supernatant was removed, and the cell pellet was washed with sterile water. Cell pellet was frozen in liquid nitrogen and stored at −80°C. Then, the cells were resuspended in 300 μl of lysis buffer (200 mM lithium acetate and 1% SDS) and incubated at 95°C for 2 minutes. Next, cells were frozen in liquid nitrogen for 2 minutes and subsequently heated at 95°C for 1 minute. This procedure was repeated twice. Samples were vortexed, and 300 μl chloroform was added and vortexed again. Samples were centrifuged at 20,000xg for 3 minutes. The upper phase was transferred to a new tube and genomic DNA was precipitated using 96% ethanol overnight at −20°C. Next, samples were centrifuged at 20,000xg for 15 minutes at 4°C. Pellets were dried at room temperature and diluted in Tris-EDTA buffer (TE) buffer (cat. no 93283, Sigma-Aldrich). One nanogram of DNA was used for quantitative PCR. Primers for nuclear and mitochondrial DNA genes are listed in **Supplementary Table 5**. The experimental amplification factor of each primer pair was used to calculate the linear form expression values of each gene. The mtDNA copy number was calculated by dividing the geometric mean expression of mitochondrial DNA genes (*COX2, COX3*) by the geometric mean expression of nuclear DNA genes (*ACT1, MIP1, MRX6*).

#### Preparation of protein extracts and western blot analysis

Four OD600 units of yeast cells were collected and resuspended in 1 ml of ddH2O, 148 μl of 2 M NaOH, 12 μl 100% beta-mercaptoethanol and 13 μl 100 mM PMSF. Samples were vortexed and incubated on ice for 10 minutes. Next, 50% of TCA was added, and incubated for another 20 minutes on ice. Samples were centrifuged for 10 minutes at 20,000xg at 4°C. Supernatants were removed, and protein pellets were washed with 100% ice-cold acetone. After centrifugation, protein pellets were dried at room temperature and resuspended in 2x Laemmli sample buffer with 50 mM DTT. Next, samples were incubated at 95°C for 5 minutes. Samples were resolved by SDS-PAGE (running buffer composition 25 mM Tris, 192 mM glycine, 0.1% SDS) using the indicated percentage of the gel. Transfer was performed on PVDF membrane (0.45 μm) using VWR® PerfectBlue, semi-dry electroblotters. Following commercially available antibodies were used: anti-Pgk1 (1:2000; cat. no 459250; Thermo Fisher Scientific), anti-actin (Act1) (1:1000; cat. no MAB1501; Sigma-Aldrich), anti-lipoic acid (1:2000; cat. no 437695-100UL; Sigma-Aldrich), anti-rabbit IgG secondary antibody (1:10,000; cat. no A9169; Sigma-Aldrich), anti-mouse IgG secondary antibody (1:10,000; cat. no A4416-1ML; Sigma-Aldrich). All other primary antibodies were custom-generated in rabbits and used at a concentration of 1:500 in 5% non-fat milk in 1x TBS. Chemiluminescence protein signals were detected with X-ray films. Images were digitally processed using Photoshop software. Densitometry measurements were performed to quantify signals from the western blot analysis using ImageJ 1.53t software (National Institutes of Health, Bethesda, MD, USA).

#### Analysis of Puf3-HA phosphorylation

HA-tagged PUF3 yeast strains were grown either in liquid YPD medium (composed of 1% yeast extract, 1% Bacto peptone, and 2% glucose) YPG (composed of 1% yeast extract, 1% Bacto peptone, and 3% glycerol) with shaking at 150 rpm at 28°C. To obtain cell extracts, the yeast cells were grown to OD600 of 0.7 – 0.8 and harvested by centrifugation at 1000×g for 3 minutes at room temperature (RT). Pelleted cells were suspended in RIPA lysis buffer (50 mM Tris-HCl pH 7.5, 150 mM NaCl, 0.1% SDS, 1% Igepal CA-630) supplemented with 1x complete Mini EDTA-free Protease Inhibitor Cocktail (cat. no 05892791001, Merck, Roche) and disrupted by vigorous shaking with glass beads (cat. no G8772, Merck, Sigma-Aldrich) followed by centrifugation at 15,000×g for 20 minutes at 4°C. The protein concentration in the supernatant was determined by the Bradford procedure using Bio-Rad Protein Assay Dye Reagent Concentrate (cat. no 5000006, Bio-Rad) and bovine serum albumin (BSA, cat. no A4503, Merck, Sigma-Aldrich) used for preparation of the standard curve. Protein samples (20 μg of total cell extracts) were resolved by SDS-PAGE. Phos-tag gels: 8% polyacrylamide supplemented with 50 μM Phos-tag (cat. no AAL-107, Wako Chemicals) and 100 μM ZnCl2 were run with the constant current of 25 mV for 5 h at 4°C. Dephosphorylated samples were used as controls; for dephosphorylation, 20 μl of total cell extracts were incubated for 1 hours at 37°C with 10 U of FastAP Thermosensitive Alkaline Phosphatase – FAP (cat. no EF065, ThermoFisher Scientific). Finally, all samples were diluted in SDS-PAGE loading buffer and heated for 5 minutes at 95 °C. After electrophoresis, the gels were washed twice for 10 minutes with transfer buffer (25 mM Tris, 236 mM glycine, 20% methanol) supplemented with 10 mM EDTA pH 8.0. The transfer of proteins to nitrocelulose membranes (Amersham Protran 0.45µm NC, cat. no 10600002, Cytiva) was carried out with a constant current of 30 mV for overnight using wet-transfer. Protein immunodetection was performed using specific primary antibodies anti-HA (cat. no 9658, Sigma-Aldrich) or anti-GAPDH (cat. no 60004-1-Ig, Proteintech). For visualization of the protein-IgG complex, a chemiluminescent signal was generated with the use of the Clarity Western ECL substrate (cat. no 170-5061, Bio-Rad) and acquired using the ChemiDoc MP imaging system (Bio-Rad).

#### Polysome profiling

The wild-type, *rpl40a*Δ, *rpl40b*Δ strains grown in YPD (2% glucose) medium until OD600 0.70–1.0 or in YPG (3% glycerol) medium until OD600 0.6–0.8. Before collection, cells were treated with 90 μg/ml cycloheximide (cat. no C7698, Sigma–Aldrich) for 5 minutes. Cell pellets were frozen in liquid nitrogen and stored at −80°C. Cells were lysed using a lysis buffer (10 mM Tris–HCl, pH 7.5, 100 mM NaCl, 30 mM MgCl2, 100 μg/ml cycloheximide, 1 mM PMSF, 10 ng/ml aprotinin, 10 nM leupeptin, 1 nM pepstatin A, 1x RNase inhibitor (cat. no R7397, Sigma–Aldrich), 6 mM beta-mercaptoethanol and 200 ng/ml heparin). Cells were disrupted by acid-washed glass beads (cat. no. G8772, Sigma–Aldrich) in a mini vortex mixer (Disruptor Genie, Scientific Industries Inc.) at maximum speed for 6 minutes at 4°C. The cell debris was removed through centrifugation at 15,294xg for 10 minutes at 4°C. Five absorbance units (A260) of cell lysate were loaded on a 10%-50% sucrose gradient in a thin wall polypropylene tube (cat. no 331372, Beckman Coulter, Inc.). Sucrose density gradients were freshly made using the Gradient Master 108 (BioComp Instruments, Inc.). Two sucrose solutions, 10% and 50%, were prepared in a buffer contained 50 mM Tris–HCl, pH 7.0, 50 mM NH4Cl, 12 mM MgCl2 and 1 mM DTT. Tubes were ultracentifuged at 38,000 rpm (acceleration: 5, deceleration: 1) at 4°C for 2 hours using the TH-641 rotor and the Sorvall WX 90 ultracentrifuge (Thermo Scientific). Samples were fractionated through a density gradient fractionation system (Brandel) that continuously monitored the absorbance at 254 nm.

### Isolation of mitochondria

#### For proteomics and lipidomics analysis

Yeast mitochondria were isolated from 1500 ml cultures (OD600 ∼1.1) grown in YPG. Supernatant was removed, cell pellet was resuspended in DTT buffer (100 mM Tris-SO4, pH 9.4, 10 mM DTT) and incubated at 30°C for 15 minutes at 100 rpm. After discarding the supernatant, the cell pellet was washed in Z buffer (1.2 M sorbitol, 20 mM KPi, pH 7.4). Next, the cell pellet was resuspended in Z buffer with 4.5 mg Zymolyase 20T (cat. no 120491-1, Amsbio) per 1 g of cell pellet. Cells were incubated at 30°C for 30 minutes at 100 rpm. The cell pellet was washed with Z buffer. The cell pellet was homogenized in a Potter-Elvehjem homogenizer using 6.5 ml homogenization buffer (0.6 M sorbitol, 10 mM Tris-HCl pH 7.4, 0.2% BSA, 1 mM PMSF) per 1 g of pellet. The cell pellet debris was pelleted at 1900xg for 5 minutes at 4°C. The supernatant was centrifuged at 12,000 rpm (rotor SORVALL SS-34, Thermo Scientific) for 15 minutes at 4°C. The pellet was resuspended in 1 ml of SM buffer (250 mM sucrose, 10 mM MOPS-KOH, pH 7.2).

#### For analysis of respiration and ATP synthesis

Mitochondria were prepared from yeast cells grown in YPG to an OD600 of 4 by the enzymatic method described previously ^63^. The cell pellet was resuspended in SH buffer (100 mM Tris, pH 9.3, 500 mM beta-mercaptoethanol) and incubated at 30°C for 10 minutes. Then, after centrifugation of cells, the cell pellet was washed twice in KCl buffer (10 mM Tris, pH 7.0, 500 mM KCl) and suspended in Zymolyase buffer (1.35 M sorbitol, 30 mM NaPi, pH 5.8, 10 mM citric acid, 1 mM EGTA) with 10.5 mg Zymolyase 20T per 1 g of cell pellet. After incubation at 30°C for 30 minutes, the spheroplasts were washed twice with Washing buffer (0.75 M sorbitol, 0.4 M mannitol, 10 mM Tris-maleate pH 6.8, 0.1% BSA). Then, they were suspended in 45 ml of Homogenization buffer (0.6 M mannitol, 10mM Tris-maleate pH 6.8, 0.2% BSA, 2 mM EGTA) and homogenized in a Waring Commercial Lab Blender 3 times for 5 seconds. The cell debris was pelleted at 1470xg for 8 minutes at 4°C. The supernatant was centrifuged at 12,000xg for 10 minutes at 4°C. The pellet of mitochondria was washed once with the Recuperation buffer (0.6 M mannitol, 10 mM Tris-maleate, pH 6.8, 2 mM EGTA) and resuspended in 0.3 ml of Recuperation buffer containing protease inhibitors cocktail (cat. no 04693132001, Roche).

#### Oxygen consumption, ATP synthesis, and hydrolysis activities in isolated mitochondria

Mitochondria were diluted to 0.075 g/ml in respiration buffer (10 mM Tris-maleate, pH 6.8, 0.65 M mannitol, 0.36 mM EGTA, and 5 mM Tris-phosphate). Oxygen consumption rates were measured using a Clarke electrode in the presence of 4 mM NADH (state 4 respiration), 150 µM ADP (state 3 respiration), as previously described ^64^. A higher concentration of ADP (750 µM) was used to measure the rate of ATP synthesis in the absence and presence (3 μg/ml) of oligomycin. Aliquots were taken every 15 seconds, and the production of ATP, after stopping the reaction with 3.5% (w/v) perchloric acid and 12.5 mM EDTA, was quantified using the Kinase-Glo Max Luminescence Kinase Assay (cat. no V6711, Promega) and a Beckman Coulter Paradigm Plate Reader.

#### Proteomics analysis of isolated mitochondria

Isolated mitochondria in SM buffer (250 mM sucrose, 10 mM MOPS-KOH, pH 7.2) were centrifuged 15,000xg for 2 minutes at 4°C, supernatant was discarded. Mitochondrial pellet was resuspended in 100 μl 8 M urea with 2 mM PMSF. Samples were lysed by syringe with a 22-gauge needle and incubated at 30°C for 15 minutes. Samples were centrifuged (2 minutes, 20,800xg, 20°C) and supernatants were transferred to new tubes. Protein concentration was determined using the microplate method and Pierce™ BCA Protein Assay Kit (cat. no 23225, Thermo Scientific). Sixty micrograms of protein per sample were collected into a Protein LoBind tube for in-solution digestion. Then, 4 μl of 1% ProteaseMAX™ Surfactant (cat. no V2072, Promega) and 50 mM NH4HCO3 to a final volume of 93.5 μl were added. Next, samples were reduced by adding 1 μl of 0.5 M DTT and incubated at 56°C for 20 minutes. Next, 2.7 μl of 0.55 M 2-chloroacetamide (CAA, cat. no 22790, Sigma-Aldrich) was added and incubated at room temperature in the dark for 15 minutes. In the last step, 1 μl of 1% ProteaseMAX™ Surfactant and 1.8 μl of 1 μg/μl trypsin were added to the sample. Samples were digested by trypsin for 17 hours at 37°C and 400 rpm. The digestion was stopped by adding TFA to the final pH ∼2. Peptides were desalted using Pierce^®^ C18 Tips (cat. no 87784, Thermo Scientific).

#### LC-MS/MS analysis

Before LC-MS/MS measurement, the samples were resuspended in 0.1% TFA, 2% acetonitrile in water. Chromatographic separation was performed on an Easy-Spray Acclaim PepMap column 50 cm length × 75 µm inner diameter (Thermo Fisher Scientific) at 55°C by applying 120 minutes’ acetonitrile gradients in 0.1% aqueous formic acid at a flow rate of 300 nl/minute. An UltiMate 3000 nano-LC system was coupled to a Q Exactive HF-X mass spectrometer via an Easy-Spray source (all Thermo Fisher Scientific). The Q Exactive HF-X was operated in data-dependent mode with survey scans acquired at a resolution of 120,000 at m/z 200. Up to 12 of the most abundant isotope patterns with charges 2–5 from the survey scan were selected with an isolation window of 1.3 m/z and fragmented by higher-energy collision dissociation (HCD) with normalized collision energies of 27, while the dynamic exclusion was set to 30 s. The maximum ion injection times for the survey scan and the MS/MS scans (acquired with a resolution of 15,000 at m/z 200) were 45 and 150 ms, respectively. The ion target value for MS was set to 3 × 1e6 and for MS/MS to 1e5, and the intensity threshold for MS/MS was set to 6.7e3.

#### Analysis of proteomics data

Raw data were processed using MaxQuant version 2.2.0.0 and parameters: variable modifications — oxidation (M), acetyl (protein N-term), deamidation (NQ); fixed modifications — carbamidomethyl (C); digestion: trypsin/P; Label-free quantification — LFQ (min. ratio count 2); Global parameters — *S. cerevisiae* 6727 FASTA reviewed sequences from UniprotKB (canonical and isoform), downloaded on 24th May 2023. Results were analyzed using Perseus v2.0.6.0. The dataset was filtered from potential contaminants, proteins only identified by site, and reverse database hits. Proteins identified based on at least one unique peptide were kept. LFQ intensities were log2-transformed and minimum 3 valid values in at least one group were kept. Missing values were imputed from a normal distribution. In total, 976 proteins were identified in isolated mitochondria samples. Data were normalized by median subtraction of the columns. Using built-in statistical analysis in Perseus v2.0.9.0, *p*-values between two groups (*rpl40a*Δ vs. WT; *rpl40b*Δ vs. WT) were determined by two-sided Student’s *t*-test with Benjamini-Hochberg adjustment of 0.05). The dataset was limited to the proteins annotated in Gene Onthology Cellular Component (GO CC) as mitochondrion (662 proteins). Volcano plots were generated in RStudio version 2024.04.2+764 employing the following work packages: ggplot2, dplyr, ggrepel, stringr. Enrichment analysis was conducted using ShinyGO 0.85.1 ^65^ and annotation database derived from STRING-db v12 for *Saccharomyces cerevisiae* (taxonomy ID 4932). The list of proteins annotated to GO Cellular Component “mitochondrion” were used as a background (1286 proteins). The false discovery rate (FDR) was set on 0.05. Results of pathway enrichment are in **Supplementary Table 7**.

### Fatty acid quantification by gas chromatography

#### Extraction of fatty acids from culture medium

Fatty acid profile analysis of the culture medium was performed according to the previously published method ^66^. Briefly, yeast cells were separated from the medium, and 0.2 mg heptanoic acid was added to 10 ml supernatant as an internal standard. Then, 1 ml of 1 M HCl and 2.5 ml of methanol–chloroform solution (1:1) were added. The solution was shaken for 5 minutes and centrifuged (3000xg, 10 minutes). The chloroform layer was recovered and evaporated overnight. Samples were dissolved in 200 µl of toluene, mixed with 1.5 ml of methanol and 300 µl of 8% (w/v) HCl solution, vortexed, and incubated at 100°C for 3 hours to form fatty acid methyl esters (FAMEs). Then, the samples were cooled, 1 ml H2O and 1 ml hexane were added, and shaken. The organic phase was transferred to a gas chromatography (GC) vial.

#### Extraction of fatty acids from mitochondria

Heptanoic acid was added as an internal standard. One milliliter of reagent R1 (150 g NaOH in 50% methanol) was added to the mitochondrial pellet for saponification, then the samples were vortexed, and incubated at 100°C for 30 min. After cooling down the samples, 2 ml of reagent R2 (mixture of 325 ml 6 M HCl and 275 ml methanol) was added, vortexed and incubated at 80°C for 10 min. After cooling the samples, 4 ml of reagent R3 (the mixture of hexane and methyl tert-butyl ether, 1:1) was added. Samples were mixed on a rotary stirrer for 10 min. The reagent R4 (1.2% NaOH) was added to the organic phase, and samples were stirred gently for 5 min. A few drops of saturated NaCl were added. The fatty acid extract was transferred to a gas chromatography vial.

#### GC-FID analysis of FAMEs

In both types of samples (culture media and mitochondria samples), fatty acid methyl esters were separated by the Agilent 7890A Gas Chromatograph (Agilent Technologies, USA) with the 7683 Automatic Liquid Sampler injector (Agilent Technologies, USA) and a flame ionization detector (FID, Agilent Technologies). The system was equipped with the DB-FFAP 250°C capillary column (30 m x 250 µm x 0.25 µm; Agilent Technologies, USA). The following GC column oven temperature program was applied: run time 100.5 min, start at 80°C and hold for 8 min, increase at the rate of 4°C/min to 100°C and hold for 3 min, increase at the rate of 4°C/min to 150°C and hold for 5 min, increase at the rate of 2°C/min to 175°C and hold for 5 min, increase at the rate of 2°C/min to 200°C and hold for 5 min, then increase at the rate of 2°C/min to 250°C and hold for 7 min. FAMEs were identified and quantified by comparison of their retention time with fatty acid methyl esters standards (F.A.M.E. Mix C8 - C24 standard; cat. no CRM18918; Supelco). ChemStation software (Agilent Technologies, USA) was used for data acquisition and quantification. The automatic integration of peaks was used, and a gate window integration of 0.5 minute was set. The extraction efficiency was in the range of 91% to 99%. Considering the extraction efficiency for each sample, the mean fatty acid content of three biological replicates per group was calculated.

#### Lipid extraction and lipidomics analysis

Lipid extraction was performed using an in-house protocol for untargeted lipidomics, according to the previously published method ^67^. Thawing and all the steps of sample preparation were conducted on ice. Samples were first subjected to homogenization by the addition of 225 µl of ice-cold methanol and zirconium beads. Next, the homogenization protocol was performed using PreCellys Evolution Orbital Homogenizer (Bertin Technologies) during 10 cycles of vigorous shaking at 4500 rpm for 20 seconds, 15-second rest at 4°C. Subsequently, 750 µl of ice-cold methyl-tert-butyl ether was added and samples were vortexed for 10 minutes at 1500 rpm, 8°C using ThermoMixer (Eppendorf). Phase separation was achieved by adding 188 µl of ice-cold MQ grade water and shaking the samples for an additional 60 seconds. Samples were centrifuged for 10 minutes at 14,000 rpm, 4°C, and the upper organic layer was transferred to new polypropylene tube. Finally, samples were evaporated to dryness under gentle stream of nitrogen at 50°C, after which the dry residues were reconstituted in methanol/toluene solution (9:1, v/v).

Lipids were separated using ACQUITY UPLC BEH C18 1.7 µm, 2.1 mm × 100 mm column (Waters). Mobile phase A consisted of ACN/H2O (7:3, v/v) with 10 mM ammonium acetate and mobile phase B consisted of IPA/ACN (9:1, v/v) with 10 mM ammonium acetate. The chromatographic gradient is summarized in **Supplementary Table 8.** Column oven was kept at 50°C and the flow was maintained at 0.4 ml/min.

Mass spectra were acquired separately in positive and negative mode using heated electrospray ionization source (HESI) using Data Dependent Acquisition mode (DDA). The heated capillary was kept at 380°C, sheath gas 25 [a.u.], auxiliary gas flow 10 [a.u.] and sweep gas to 1 [a.u.]. Capillary voltage was set to 4.0 kV for both ionization modes. The transfer capillary temperature was maintained at 320°C and the S-lens RF value 85.0 [a.u.]. MS1 resolution was set to 70,000 max injection time (IT) 120 ms and scan range of 120–1600 m/z. MS2 resolution was set to 35,000 max IT 90 ms and stepped collision energy 20, 30 [a.u.].

RAW files were uploaded to Compound Discoverer 3.3.3 (CD, Thermo Scientific) and analyzed using a protocol presented previously ^67,68^. Briefly, “aligned” and “unaligned” workflows were employed which consisted of subsequent steps including: Input Files, Select Spectra, Align Retention Times, Detect Compound, Group Compound, Fill Gaps, Mark Background.

Lipid identification was processed using Lipidex 1.1 (Coon Laboratories) software. RAW files were converted to MGF files using MSConvertGUI (ProteoWizard, Stanford University), who were then searched with Spectrum Searcher algorithm using built-in databases. The Aligned and Unaligned data sheets from CD software were used for final metabolite determination employing the Peak Finder algorithm. Details on specific method parameters are given in the **Supplementary Table 6**. MetaboAnalyst 6.0 ^69^ was used for data processing and statistical analysis using sample normalization by median, log2 transformation, Principal Component Analysis (PCA) and Student’s *t*-test. Unidentified lipids where not shown on the Volcano plots.

## Data availability

The authors confirm that all relevant data are included in the main manuscript and Supplementary files provided with the manuscript. The mass spectrometry proteomics data have been deposited to the ProteomeXchange Consortium via the PRIDE ^70^ partner repository with the dataset identifier PXD077150 and 10.6019/PXD077150. The mass spectrometry lipidomics data have been deposited to MetaboLights ^71^ repository with the study identifier MTBLS14310. The materials that were generated within this work are available upon request from the corresponding author (U.T.).

## Author contribution

KPL – data curation, investigation, formal analysis, methodology, validation, visualization, writing – original draft, funding acquisition; KG – investigation, formal analysis, visualization; MS, SSG, KJ – investigation, writing – review and editing; MR – data curation, methodology; RK – investigation, methodology, writing – review and editing; AW, EM – investigation; MT – investigation; writing – review and editing; MM, AS – investigation, methodology; UT – conceptualization, investigation, validation, funding acquisition, supervision, writing – original draft. All authors agreed to the final version of the manuscript.

## Acknowledgments

Proteomic measurements were performed at the Proteomics Core Facility, IMol Polish Academy of Sciences. Special thanks to Dorota Stadnik and Remigiusz Serwa (IMol PAS) for help with sample preparation and LC-MS/MS measurements. The GC-FID analysis was performed by the Laboratory of Environmental Instrumental Analyzes at the University of Warsaw. The PAMS366 plasmids and calcineurin activity methodology were kindly provided by Joanna Kamińska (Institute of Biochemistry and Biophysics PAS, Poland). The petite frequency assay methodology was kindly provided by Aneta Kaniak-Golik (Institute of Biochemistry and Biophysics PAS, Poland).

## References

1. Komili, S., Farny, N.G., Roth, F.P., and Silver, P.A. (2007). Functional specificity among ribosomal proteins regulates gene expression. Cell 131, 557–571.

2. Ghulam, M.M., Catala, M., and Abou Elela, S. (2020). Differential expression of duplicated ribosomal protein genes modifies ribosome composition in response to stress. Nucleic Acids Res 48, 1954–1968.

3. Pietras, P.J., Wasilewska-Burczyk, A., Pepłowska, K., Marczak, Ł., Tyczewska, A., and Grzywacz, K. (2024). Dynamic protein composition of Saccharomyces cerevisiae ribosomes in response to multiple stress conditions reflects alterations in translation activity. International Journal of Biological Macromolecules 268, 132004.

4. Petibon, C., Parenteau, J., Catala, M., and Elela, S.A. (2016). Introns regulate the production of ribosomal proteins by modulating splicing of duplicated ribosomal protein genes. Nucleic Acids Res 44, 3878–3891.

5. Komili, S., Farny, N.G., Roth, F.P., and Silver, P.A. (2007). Functional Specificity among Ribosomal Proteins Regulates Gene Expression. Cell 131, 557–571.

6. Ferretti, M.B., and Karbstein, K. (2019). Does functional specialization of ribosomes really exist? Rna 25, 521–538.

7. Rao, S., Peri, S., Hoffmann, J., Cai, K.Q., Harris, B., Rhodes, M., Connolly, D.C., Testa, J.R., and Wiest, D.L. (2019). RPL22L1 induction in colorectal cancer is associated with poor prognosis and 5-FU resistance. PLoS One 14, e0222392.

8. Malik Ghulam, M., Catala, M., Reulet, G., Scott, M.S., and Abou Elela, S. (2022). Duplicated ribosomal protein paralogs promote alternative translation and drug resistance. Nature Communications 13, 4938.

9. O’Leary, M.N., Schreiber, K.H., Zhang, Y., Duc, A.C., Rao, S., Hale, J.S., Academia, E.C., Shah, S.R., Morton, J.F., Holstein, C.A., et al. (2013). The ribosomal protein Rpl22 controls ribosome composition by directly repressing expression of its own paralog, Rpl22l1. PLoS Genet 9, e1003708.

10. Park, S., Fitzgerald, F., Yang, Y.M., and Karbstein, K. (2026). A deep dive into functional ribosome specialization. J Cell Biol 225.

11. Segev, N., and Gerst, J.E. (2018). Specialized ribosomes and specific ribosomal protein paralogs control translation of mitochondrial proteins. J Cell Biol 217, 117–126.

12. Zou, Q., and Qi, H. (2021). Deletion of ribosomal paralogs Rpl39 and Rpl39l compromises cell proliferation via protein synthesis and mitochondrial activity. The International Journal of Biochemistry & Cell Biology 139, 106070.

13. de Alteriis, E., Carteni, F., Parascandola, P., Serpa, J., and Mazzoleni, S. (2018). Revisiting the Crabtree/Warburg effect in a dynamic perspective: a fitness advantage against sugar-induced cell death. Cell Cycle 17, 688–701.

14. Tyczewska, A., and Bąkowska-Żywicka, K. (2025). Stress-induced ribosomal heterogeneity in Saccharomyces cerevisiae: from protein paralogs to regulatory noncoding RNAs. FEMS Yeast Res 25.

15. Sun, M., Shen, B., Li, W., Samir, P., Browne, C.M., Link, A.J., and Frank, J. (2021). A Time-Resolved Cryo-EM Study of Saccharomyces cerevisiae 80S Ribosome Protein Composition in Response to a Change in Carbon Source. Proteomics 21, e2000125.

16. Fernández-Pevida, A., Rodríguez-Galán, O., Díaz-Quintana, A., Kressler, D., and de la Cruz, J. (2012). Yeast ribosomal protein L40 assembles late into precursor 60 S ribosomes and is required for their cytoplasmic maturation. J Biol Chem 287, 38390–38407.

17. Ma, C., Wu, S., Li, N., Chen, Y., Yan, K., Li, Z., Zheng, L., Lei, J., Woolford, J.L., Jr., and Gao, N. (2017). Structural snapshot of cytoplasmic pre-60S ribosomal particles bound by Nmd3, Lsg1, Tif6 and Reh1. Nat Struct Mol Biol 24, 214–220.

18. Wang, X., and Chen, X.J. (2015). A cytosolic network suppressing mitochondria-mediated proteostatic stress and cell death. Nature 524, 481–484.

19. Tiwari, S., Singh, A., Gupta, P., K, A., and Singh, S. (2023). UBA52 Attunes VDAC1-Mediated Mitochondrial Dysfunction and Dopaminergic Neuronal Death. ACS Chemical Neuroscience 14, 839–850.

20. Ma, S., Liu, Q., Han, W., Liu, Z., Jin, S., Wu, H., and Hua, W. (2026). UBA52 Overexpression Ameliorates Intracerebral Hemorrhage-Associated Neuronal Apoptosis and Mitochondrial Dysfunction: A Protective Role in Neurons. Molecular Neurobiology 63, 371.

21. Lee, A.S., Burdeinick-Kerr, R., and Whelan, S.P. (2013). A ribosome-specialized translation initiation pathway is required for cap-dependent translation of vesicular stomatitis virus mRNAs. Proc Natl Acad Sci U S A 110, 324–329.

22. Tsai, H.Y., Liu, L., Fleming, R.H., Mintseris, J., Ekanayake, D., Gu, X., Gygi, S.P., and Lee, A.S.Y. (2025). Ribosome remodeling drives translation adaptation during viral infection and cellular stress. bioRxiv.

23. Lee, S.-O., Kelliher, J.L., Song, W., Tengler, K., Sarkar, A., Dray, E., and Leung, J.W.C. (2023). UBA80 and UBA52 fine-tune RNF168-dependent histone ubiquitination and DNA repair. Journal of Biological Chemistry 299, 105043.

24. Topf, U., Suppanz, I., Samluk, L., Wrobel, L., Boser, A., Sakowska, P., Knapp, B., Pietrzyk, M.K., Chacinska, A., and Warscheid, B. (2018). Quantitative proteomics identifies redox switches for global translation modulation by mitochondrially produced reactive oxygen species. Nat Commun 9, 324.

25. Houtkooper, R.H., Mouchiroud, L., Ryu, D., Moullan, N., Katsyuba, E., Knott, G., Williams, R.W., and Auwerx, J. (2013). Mitonuclear protein imbalance as a conserved longevity mechanism. Nature 497, 451–457.

26. Liu, Z., and Butow, R.A. (2006). Mitochondrial retrograde signaling. Annu Rev Genet 40, 159–185.

27. Jacquemyn, J., Cascalho, A., and Goodchild, R.E. (2017). The ins and outs of endoplasmic reticulum-controlled lipid biosynthesis. EMBO Rep 18, 1905–1921.

28. English, A.R., and Voeltz, G.K. (2013). Endoplasmic reticulum structure and interconnections with other organelles. Cold Spring Harb Perspect Biol 5, a013227.

29. Zhao, F., Yang, J., Li, J., Li, Z., Lin, Y., Zheng, S., Liang, S., and Han, S. (2020). Multiple cellular responses guarantee yeast survival in presence of the cell membrane/wall interfering agent sodium dodecyl sulfate. Biochem Biophys Res Commun 527, 276–282.

30. Suda, K., Moriyama, Y., Razali, N., Chiu, Y., Masukagami, Y., Nishimura, K., Barbee, H., Takase, H., Sugiyama, S., Yamazaki, Y., et al. (2024). Plasma membrane damage limits replicative lifespan in yeast and induces premature senescence in human fibroblasts. Nat Aging 4, 319–335.

31. Kingsbury, T.J., and Cunningham, K.W. (2000). A conserved family of calcineurin regulators. Genes Dev 14, 1595–1604.

32. Yamamuro, T., Katoh, D., Silva, G.M., Nishida, H., Oikawa, S., Higuchi, Y., Wang, D., Fujimoto, M., Yoshida, N., Li, M., et al. (2026). Mitochondrial control of glycerolipid synthesis by a PEP shuttle. Cell 189, 2988–3003 e2923.

33. Lapointe, C.P., Stefely, J.A., Jochem, A., Hutchins, P.D., Wilson, G.M., Kwiecien, N.W., Coon, J.J., Wickens, M., and Pagliarini, D.J. (2018). Multi-omics Reveal Specific Targets of the RNA-Binding Protein Puf3p and Its Orchestration of Mitochondrial Biogenesis. Cell Syst 6, 125–135 e126.

34. Van Vranken, J.G., Nowinski, S.M., Clowers, K.J., Jeong, M.Y., Ouyang, Y., Berg, J.A., Gygi, J.P., Gygi, S.P., Winge, D.R., and Rutter, J. (2018). ACP Acylation Is an Acetyl-CoA-Dependent Modification Required for Electron Transport Chain Assembly. Mol Cell 71, 567–580 e564.

35. de Kroon, A.I. (2007). Metabolism of phosphatidylcholine and its implications for lipid acyl chain composition in Saccharomyces cerevisiae. Biochim Biophys Acta 1771, 343–352.

36. Zhao, Q., and Sarinay Cenik, E. (2025). Is mitochondrial function at the heart of ribosome-related diseases? Trends Cell Biol 35, 815–818.

37. Lewis, A.G., Caldwell, R., Rogers, J.V., Ingaramo, M., Wang, R.Y., Soifer, I., Hendrickson, D.G., McIsaac, R.S., Botstein, D., and Gibney, P.A. (2021). Loss of major nutrient sensing and signaling pathways suppresses starvation lethality in electron transport chain mutants. Mol Biol Cell 32, ar39.

38. Wullschleger, S., Loewith, R., and Hall, M.N. (2006). TOR signaling in growth and metabolism. Cell 124, 471–484.

39. Holcik, M., and Sonenberg, N. (2005). Translational control in stress and apoptosis. Nat Rev Mol Cell Biol 6, 318–327.

40. Harding, H.P., Zhang, Y., Zeng, H., Novoa, I., Lu, P.D., Calfon, M., Sadri, N., Yun, C., Popko, B., Paules, R., et al. (2003). An integrated stress response regulates amino acid metabolism and resistance to oxidative stress. Mol Cell 11, 619–633.

41. Di Bartolomeo, F., Malina, C., Campbell, K., Mormino, M., Fuchs, J., Vorontsov, E., Gustafsson, C.M., and Nielsen, J. (2020). Absolute yeast mitochondrial proteome quantification reveals trade-off between biosynthesis and energy generation during diauxic shift. Proc Natl Acad Sci U S A 117, 7524–7535.

42. Couvillion, M.T., Soto, I.C., Shipkovenska, G., and Churchman, L.S. (2016). Synchronized mitochondrial and cytosolic translation programs. Nature 533, 499–503.

43. Chen, W., Zhao, H., and Li, Y. (2023). Mitochondrial dynamics in health and disease: mechanisms and potential targets. Signal Transduct Target Ther 8, 333.

44. Giacomello, M., Pyakurel, A., Glytsou, C., and Scorrano, L. (2020). The cell biology of mitochondrial membrane dynamics. Nat Rev Mol Cell Biol 21, 204–224.

45. Mårtensson, C.U., Doan, K.N., and Becker, T. (2017). Effects of lipids on mitochondrial functions. Biochimica et Biophysica Acta (BBA) - Molecular and Cell Biology of Lipids 1862, 102–113.

46. Baker, Z.N., Zhu, Y., Guerra, R.M., Smith, A.J., Arra, A., Serrano, L.R., Overmyer, K.A., Mukherji, S., Craig, E.A., Coon, J.J., and Pagliarini, D.J. (2025). Triacylglycerol mobilization underpins mitochondrial stress recovery. Nat Cell Biol 27, 298–308.

47. Olzmann, J.A., and Carvalho, P. (2019). Dynamics and functions of lipid droplets. Nat Rev Mol Cell Biol 20, 137–155.

48. Mathiowetz, A.J., and Olzmann, J.A. (2024). Lipid droplets and cellular lipid flux. Nat Cell Biol 26, 331–345.

49. Henne, W.M., Reese, M.L., and Goodman, J.M. (2018). The assembly of lipid droplets and their roles in challenged cells. Embo j 37.

50. Hariri, H., Rogers, S., Ugrankar, R., Liu, Y.L., Feathers, J.R., and Henne, W.M. (2018). Lipid droplet biogenesis is spatially coordinated at ER-vacuole contacts under nutritional stress. EMBO Rep 19, 57–72.

51. Seo, A.Y., Lau, P.W., Feliciano, D., Sengupta, P., Gros, M.A.L., Cinquin, B., Larabell, C.A., and Lippincott-Schwartz, J. (2017). AMPK and vacuole-associated Atg14p orchestrate μ-lipophagy for energy production and long-term survival under glucose starvation. Elife 6.

52. Contamine, V., and Picard, M. (2000). Maintenance and integrity of the mitochondrial genome: a plethora of nuclear genes in the budding yeast. Microbiol Mol Biol Rev 64, 281–315.

53. Luévano-Martínez, L.A., Forni, M.F., dos Santos, V.T., Souza-Pinto, N.C., and Kowaltowski, A.J. (2015). Cardiolipin is a key determinant for mtDNA stability and segregation during mitochondrial stress. Biochimica et Biophysica Acta (BBA) - Bioenergetics 1847, 587–598.

54. Stenberg, S., Li, J., Gjuvsland, A.B., Persson, K., Demitz-Helin, E., González Peña, C., Yue, J.-X., Gilchrist, C., Ärengård, T., Ghiaci, P., et al. (2022). Genetically controlled mtDNA deletions prevent ROS damage by arresting oxidative phosphorylation. eLife 11, e76095.

55. Ogur, M., St. John, R., and Nagai, S. (1957). Tetrazolium overlay technique for population studies of respiration deficiency in yeast. Science 125, 928–929.

56. Hess, D.C., Myers, C.L., Huttenhower, C., Hibbs, M.A., Hayes, A.P., Paw, J., Clore, J.J., Mendoza, R.M., Luis, B.S., Nislow, C., et al. (2009). Computationally driven, quantitative experiments discover genes required for mitochondrial biogenesis. PLoS Genet 5, e1000407.

57. Gościńska, K., Shahmoradi Ghahe, S., Domogała, S., and Topf, U. (2020). Eukaryotic Elongation Factor 3 Protects Saccharomyces cerevisiae Yeast from Oxidative Stress. Genes (Basel) 11.

58. Ye, J., Coulouris, G., Zaretskaya, I., Cutcutache, I., Rozen, S., and Madden, T.L. (2012). Primer-BLAST: A tool to design target-specific primers for polymerase chain reaction. BMC Bioinformatics 13, 134.

59. Damgaard, M.V., and Treebak, J.T. (2022). Protocol for qPCR analysis that corrects for cDNA amplification efficiency. STAR Protoc 3, 101515.

60. Valente, A.J., Maddalena, L.A., Robb, E.L., Moradi, F., and Stuart, J.A. (2017). A simple ImageJ macro tool for analyzing mitochondrial network morphology in mammalian cell culture. Acta Histochemica 119, 315–326.

61. Kaminska, J., Wysocka-Kapcinska, M., Smaczynska-de Rooij, I., Rytka, J., and Zoladek, T. (2005). Pan1p, an actin cytoskeleton-associated protein, is required for growth of yeast on oleate medium. Exp Cell Res 310, 482–492.

62. Tahmaz, I., Shahmoradi Ghahe, S., Stasiak, M., Liput, K.P., Jonak, K., and Topf, U. (2023). Prefoldin 2 contributes to mitochondrial morphology and function. BMC Biol 21, 193.

63. Guerin, B., Labbe, P., and Somlo, M. (1979). Preparation of yeast mitochondria (Saccharomyces cerevisiae) with good P/O and respiratory control ratios. Methods Enzymol 55, 149–159.

64. Rigoulet, M., and Guerin, B. (1979). Phosphate transport and ATP synthesis in yeast mitochondria: effect of a new inhibitor: the tribenzylphosphate. FEBS Lett 102, 18–22.

65. Ge, S.X., Jung, D., and Yao, R. (2020). ShinyGO: a graphical gene-set enrichment tool for animals and plants. Bioinformatics 36, 2628–2629.

66. Henritzi, S., Fischer, M., Grininger, M., Oreb, M., and Boles, E. (2018). An engineered fatty acid synthase combined with a carboxylic acid reductase enables de novo production of 1-octanol in Saccharomyces cerevisiae. Biotechnol Biofuels 11, 150.

67. Linke, V., Chodkowski, M., Kaszuba, K., Radkiewicz, M., Schrader, T.A., Das, H., Rana, V., Stadnik, D., Dadlez, M., Warscheid, B., et al. (2025). Integrated proteome and lipidome analyses place OCIAD1 at the mitochondria-peroxisome intersection balancing lipid metabolism. J Cell Sci 138.

68. Linke, V., Overmyer, K.A., Miller, I.J., Brademan, D.R., Hutchins, P.D., Trujillo, E.A., Reddy, T.R., Russell, J.D., Cushing, E.M., Schueler, K.L., et al. (2020). A large-scale genome-lipid association map guides lipid identification. Nat Metab 2, 1149–1162.

69. Pang, Z., Lu, Y., Zhou, G., Hui, F., Xu, L., Viau, C., Spigelman, Aliya F., MacDonald, Patrick E., Wishart, David S., Li, S., and Xia, J. (2024). MetaboAnalyst 6.0: towards a unified platform for metabolomics data processing, analysis and interpretation. Nucleic Acids Research 52, W398–W406.

70. Perez-Riverol, Y., Bai, J., Bandla, C., Garcia-Seisdedos, D., Hewapathirana, S., Kamatchinathan, S., Kundu, D.J., Prakash, A., Frericks-Zipper, A., Eisenacher, M., et al. (2022). The PRIDE database resources in 2022: a hub for mass spectrometry-based proteomics evidences. Nucleic Acids Res 50, D543–D552.

71. Yurekten, O., Payne, T., Tejera, N., Amaladoss, F.X., Martin, C., Williams, M., and O’Donovan, C. (2024). MetaboLights: open data repository for metabolomics. Nucleic Acids Res 52, D640–D646.

