## Supplementary material for "Rpl40/eL40 ribosomal protein paralogs couple cytosolic translation to mitochondrial proteome and lipid homeostasis": Document S1

### SUPPLEMENTARY FILES

### CONTENT

**Supplementary Figure 1 (Related to Figure 1).** Characterization of translation upon deletion of *RPL40* paralogs

**Supplementary Figure 2 (Related to figure 2).** Survival and mitochondrial shape

**Supplementary Figure 3 (Related to Figure 3).** Proteomics analysis of isolated mitochondria

**Supplementary Figure 4.** Analysis of mitochondrial fatty acid synthesis pathway and protein lipoylation

**Supplementary Figure 5 (Related to Figure 7).** Lipidomics analysis in total cell extracts

**Supplementary Figure 6 (Related to Figure 7).** Lipidomics analysis in mitochondrial fraction

**Supplementary Table 1.** Proteomics analysis of isolated mitochondria from WT, *rpl40a*Δ, and *rpl40b*Δ strains grown in respiratory medium at 28°C

**Supplementary Table 2.** Quantification of fatty acids abundance using gas chromatography

**Supplementary Table 3.** Lipidomic profiling of total cell extracts and isolated mitochondria from WT, *rpl40a*Δ, and *rpl40b*Δ strains grown on respiratory medium at 28°C

**Supplementary Table 4.** Yeast strains used in this study

**Supplementary Table 5.** The sequence of forward and reverse primers used in this study

**Supplementary Table 6.** Parameters for processing and analysis of lipidomics data

**Supplementary Table 7.** Enrichment analysis of GO Cellular Component terms enriched in upregulated and downregulated proteins

**Supplementary Table 8.** Summary of chromatographic binary gradient steps.

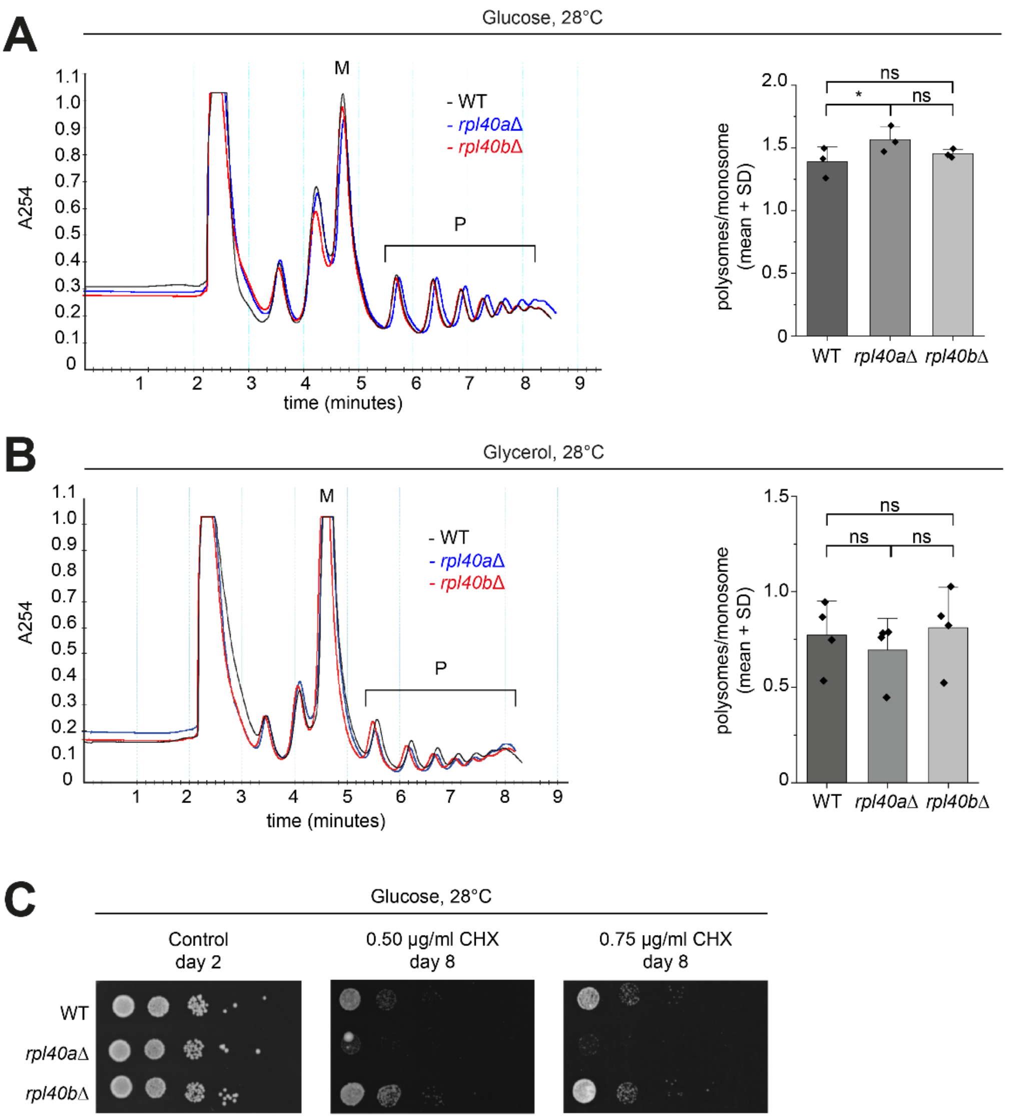


**Supplementary Figure 1 (Related to Figure 1). Characterization of translation upon deletion of *RPL40* paralogs. A, B.** Polysome profiles of wild-type (black), *rpl40a*Δ (blue) and *rpl40b*Δ (red) deletion strains grown in YPD (2% glucose) medium until $OD_{600}$ 0.70–1.0 or in YPG (3% glycerol) medium until $OD_{600}$ 0.56–0.75. Before collection, cells were treated with 90 µg/ml cycloheximide for 5 min. Five absorbance units ($A_{260}$) of cell lysate were loaded on a sucrose gradient 10%-50%, ultracentrifuged and fractionated by measuring the absorbance at 254 nm. Bar charts show the mean polysomes (P) to monosome (M) ratio and standard deviation. Significance was determined by a paired sample t test, ns $p > 0.05$, *$p \leq 0.05$, $n$ = 3–4, WT, wild-type cells. **C.** One $OD_{600}$ of wild-type and deletion strains cells were used to prepared 10-fold serial dilutions in sterile water. Two microliters of each dilution was spotted onto solid YPD (2% glucose, 2% agar) plates with 0.50 µg/ml or 0.75 µg/ml cycloheximide (CHX). Plates were incubated at 28°C for two or eight days, respectively. Representative image of three independent experiments.

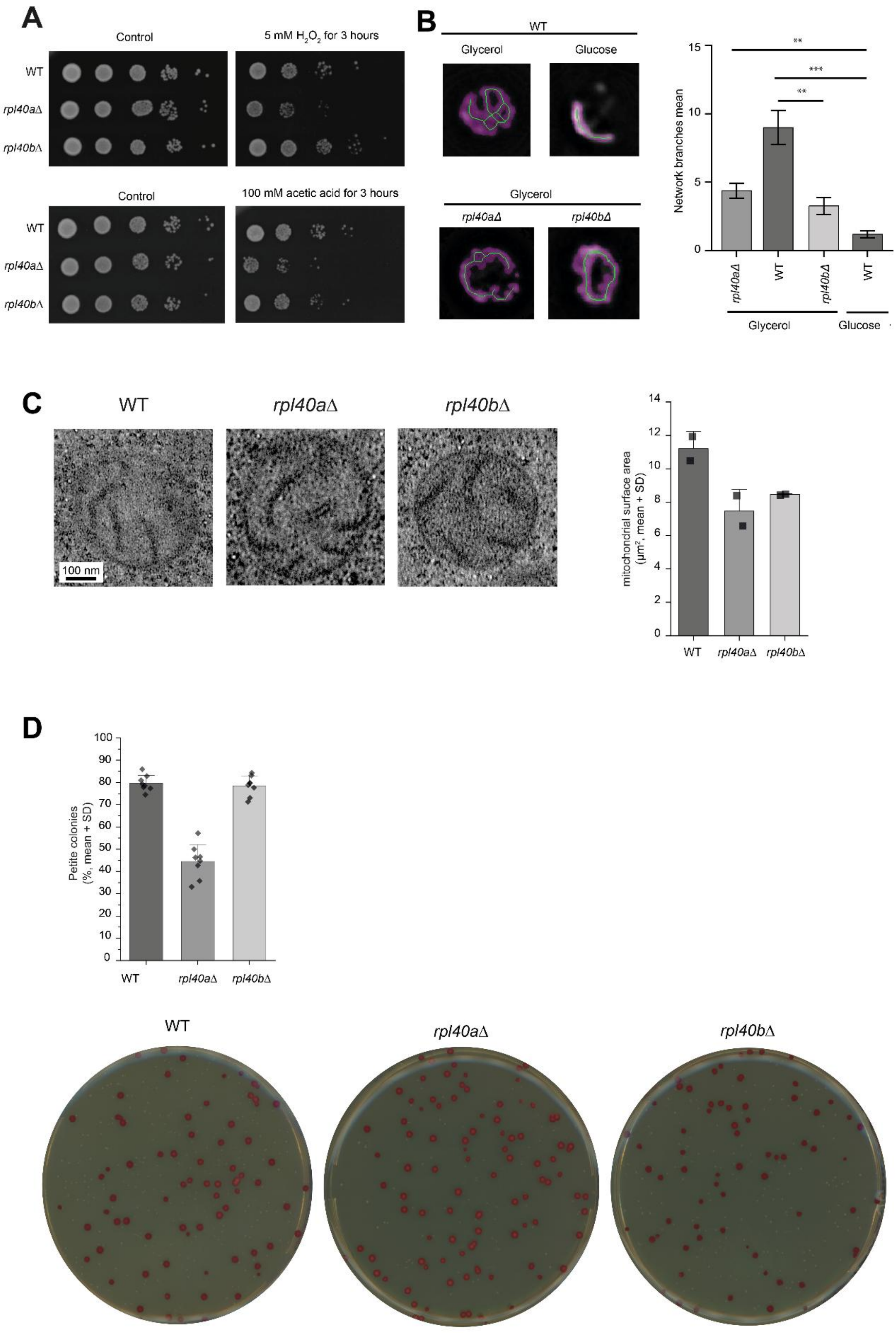


**Supplementary Figure 2 (Related to Figure 2). Survival and mitochondrial morphology. A.** Cells were grown in glucose-containing medium to the logarithmic growth phase. One $OD_{600}$ unit was pelleted, washed, and resuspended in sterile water (Control), 5 mM $H_2O_2$, or 100 mM acetic acid diluted in sterile $ddH_2O$. Cells were incubated for 3 hours at 28°C with shaking, pelleted, washed with sterile water, and 10-fold serial dilutions were prepared. Two microliters of each dilution was spotted onto solid YPD (2% glucose, 2% agar) plates. Plates were incubated for 2 days at 28°C. Representative images of 3–4 independent replicates. WT, wild-type cells **B.** (*Left*) Representative morphology of the mitochondrial network branches. Yeast cells were grown on non-fermentative medium to the logarithmic growth phase. Mitochondria were visualized using the expression of mitochondria-targeted green fluorescent protein (GFP) under a fluorescence microscope. (*Right*) Quantification of network branches represents two biological replicates and at least 9 individual cells. Data are presented as mean ± SEM, $^{**}p \leq 0.01$, $^{***}p \leq 0.001$. WT, wild-type cells. **C.** (*Left*) Representative transmission electron microscopy (TEM) images of mitochondrial cross sections from wild-type, *rpl40a*Δ and *rpl40b*Δ strains cultured in glycerol-containing medium until $OD_{600}$ 0.6–0.7. Scale bar 100 nm. (*Right*) Quantification of mitochondria area per 100 $\mu m^2$ of the cell from two biological replicates and at least 31 individual cells. Data are presented as mean + SD. $n = 2$. WT, wild-type cells. **D.** *(Upper panel)* Petite frequency assay calculated as a percentage of the number of white (petite) colonies to the total number of colonies on a plate (3% glycerol, 0.05% glucose). Data are presented as the mean of two biological replicates, where each replicate tested four colonies + SD. *(Lower panel)* Representative images of the petite frequency assay after tetrazolium overlay. Red colonies indicate respiratory competent cells, white colonies are respiratory deficient cells (petite). WT, wild-type cells.

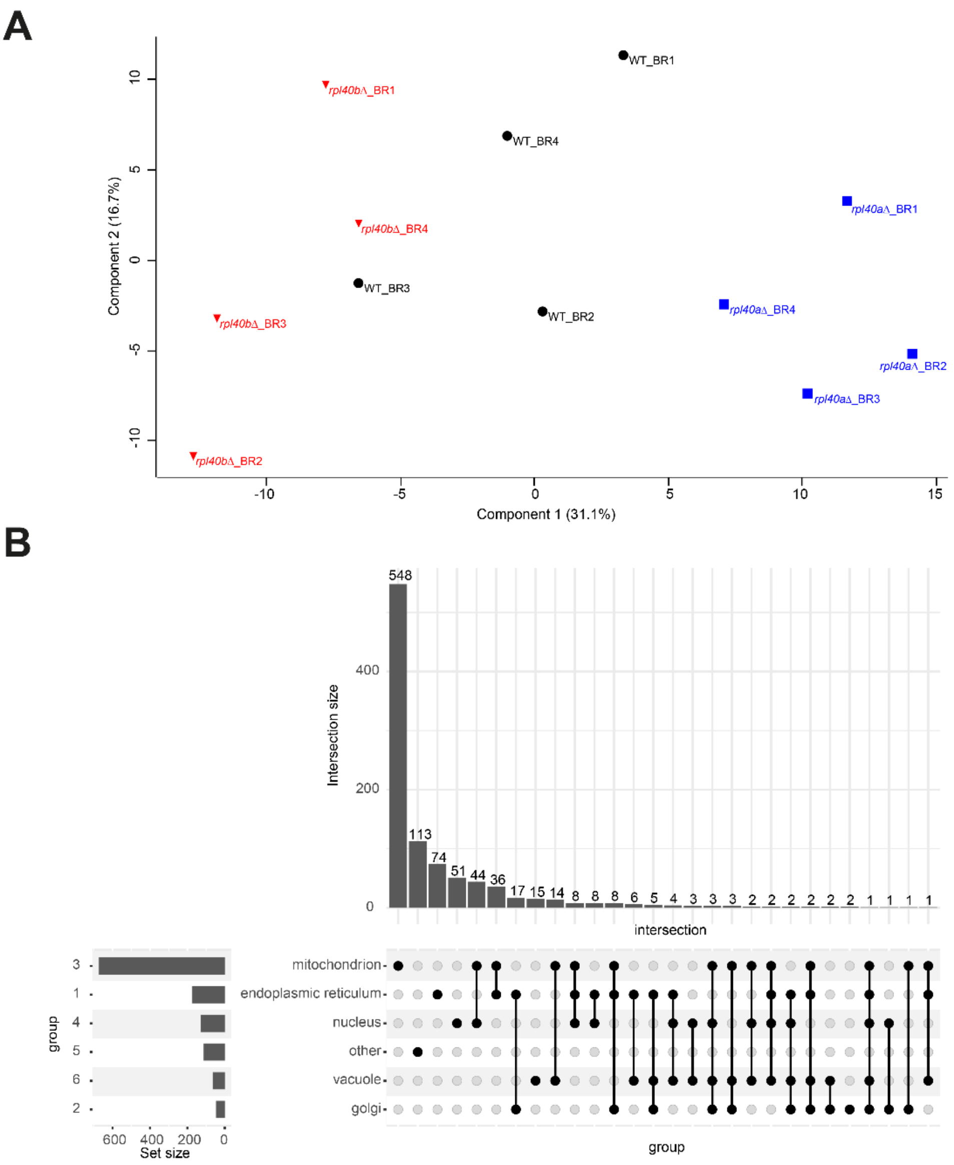


**Supplementary Figure 3 (Related to Figure 3). Proteomics analysis of isolated mitochondria. A.** Principal component analysis (PCA) of proteomes of the individual replicates of wild-type, *rpl40a*Δ and *rpl40b*Δ strains. Each strain had four independent biological replicates. Variance is indicated in brackets. **B.** Upset plot showing differential localization according to the GO annotation of all proteins identified in proteomics analysis.

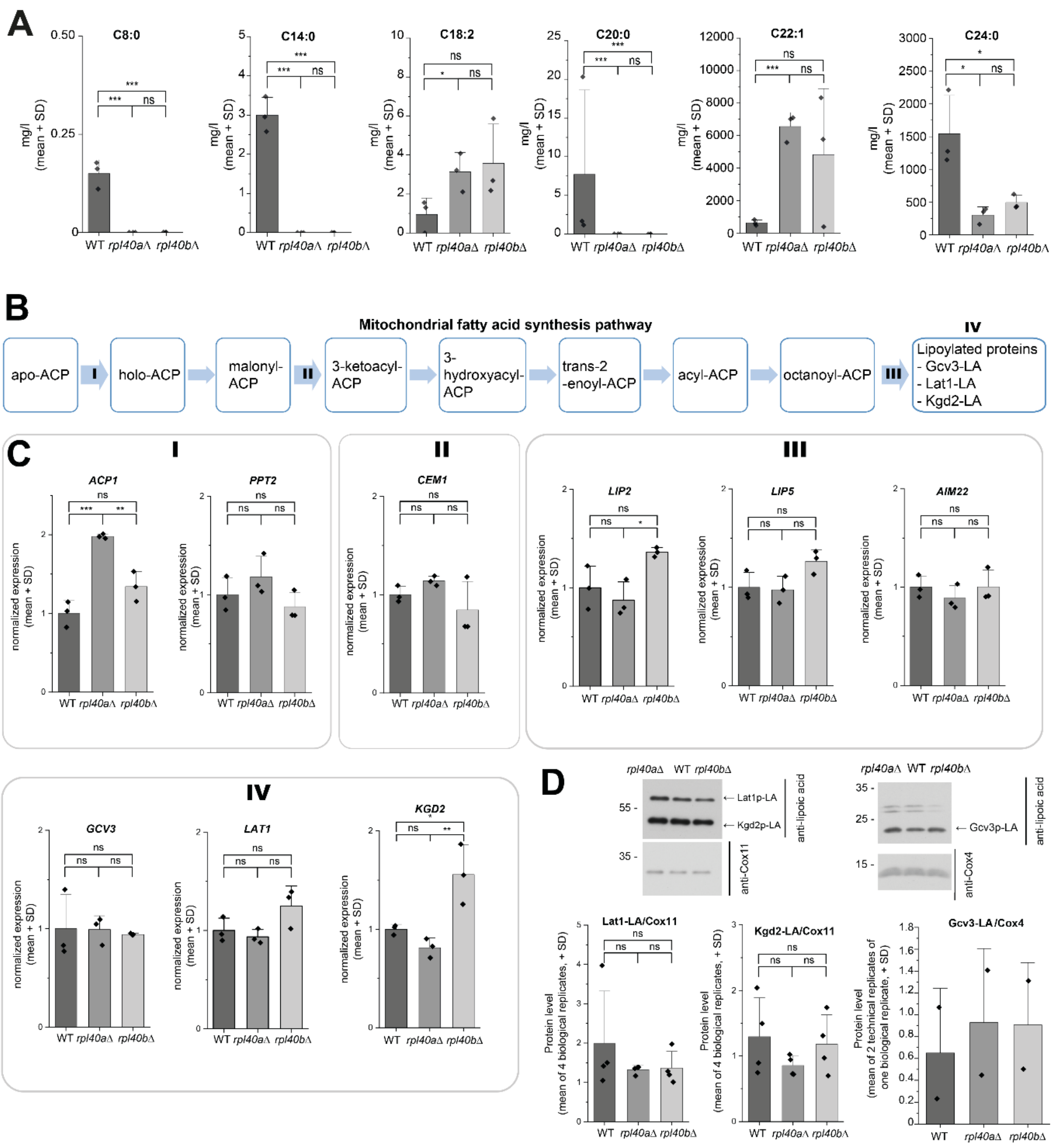


**Supplementary Figure 4. Analysis of mitochondrial fatty acid synthesis pathway and protein lipoylation. A.** Mitochondrial fractions were isolated from cultures grown on YPG (3% glycerol) medium until $OD_{600}$ 0.8–1.1. Fatty acids were extracted, separated, and analysed using gas chromatography with a flame ionization detector (GC-FID). Fatty acid methyl esters (FAMEs) were identified and quantified using an external FAME standard. For normally distributed measurements (C18:2, C22:1, C24:0) within a strain, the two-sample t-test was calculated. For not normally distributed measurements (C8:0, C14:0, C20:0) within a strain, the Kruskal-Wallis ANOVA and Conover's test were used. The graphs present the mean of three biological replicates and the standard deviation. *$p \leq 0.05$, ***$p \leq 0.001$, ns $p > 0.05$, $n$ = 3, WT, wild-type cells. **B.** Scheme of mitochondrial fatty acid biosynthesis pathway. Indicated are steps I-IV, which are analysed by qPCR shown in panel C. **C.** (I-IV) Quantitative PCR of mitochondrial fatty acid synthesis pathway genes. Wild-type (WT), *rpl40a*Δ and *rpl40b*Δ strains were grown at 28°C on YPG (3% glycerol) medium. Gene expression levels were relative to the geometric mean of the *ACT1*, *ALG9* and *TDH1*. Each measurement was normalized to the average gene expression of the wild-type strain. The graphs represent the mean and standard deviation of three biological replicates. Significance was determined by one-way ANOVA and Tukey test, ns $p > 0.05$, *$p \leq 0.05$, **$p \leq 0.01$, ***$p \leq 0.001$, $n = 3$. **D.** (*Upper panel*) Protein extracts of isolated mitochondria were separated on SDS-PAGE and analysed by western blot using the antibody against lipoic acid. (*Lower panel*) Quantification of lipoate-binding proteins (Lat1, Kgd2, Gcv3) normalized to the Cox11 and Cox4 proteins. Data are presented as mean + SD, $n$ = 1–4. Significance was determined by a paired sample *t*-test. ns, not significant. WT, wild-type cells**.**

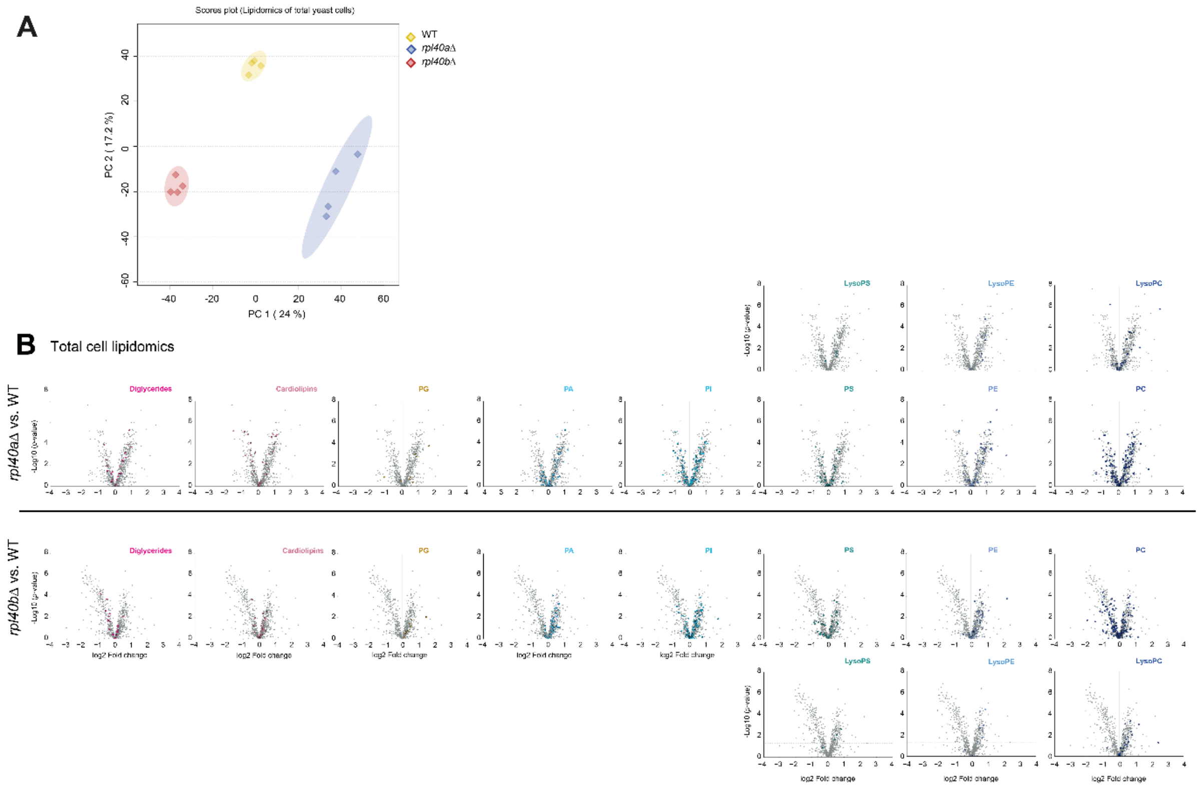


**Supplementary Figure 5 (Related to Figure 7). Lipidomics analysis of total cell extracts. A.** Principal Component Analysis (PCA) of total lipidomes of the individual biological replicates (n=4) of wild-type (WT; yellow), *rpl40a*Δ (blue), and *rpl40b*Δ (red) strains. Variance of two components (PC1 and PC2) is indicated in brackets. **B.** (*Upper panel*) Volcano plots of total cell lipidomics presenting the $\log_2$ fold change of abundance of all quantified molecular species of diglycerides, cardiolipin, phosphatidylglycerol (PG), phosphatidic acid (PA), phosphatidylinositol (PI), phosphatidylserine (PS), lysophosphatidylserine (LysoPS), phosphatidylethanolamine (PE), lysophosphatidylethanolamine (LysoPE), phosphatidylcholine (PC) and lysophosphatidylcholine (LysoPC) between *rpl40a*Δ and wild-type (WT) cells. (*Lower panel*) Volcano plots of total cell lipidomics presenting the $\log_2$ fold change of abundance of all quantified molecular species of diglycerides, cardiolipin, PG, PA, PI, PS, LysoPS, PE, LysoPE, PC and LysoPC between *rpl40b*Δ and WT cells; *n* = 4.

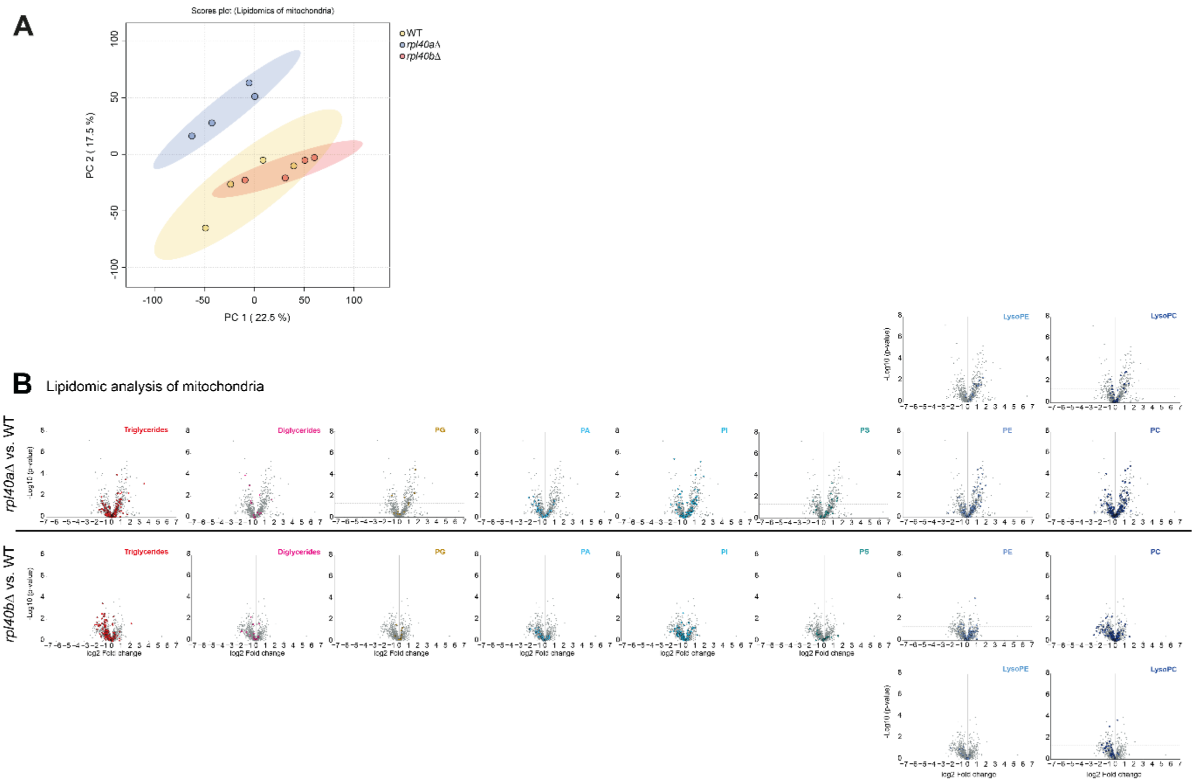


**Supplementary Figure 6 (Related to Figure 7). Lipidomics analysis of the mitochondrial fraction. A.** Principal Component Analysis (PCA) of mitochondrial lipidomes of the four individual biological replicates of wild-type (WT; yellow), *rpl40a*Δ (blue), and *rpl40b*Δ (red) strains. Variance of two components (PC1 and PC2) is indicated in brackets. **B.** (*Upper panel*) Volcano plots of mitochondria lipidomics presenting the $\log_2$ fold change of abundance of all quantified molecular species of triglycerides, diglycerides, phosphatidylglycerol (PG), phosphatidic acid (PA), phosphatidylinositol (PI), phosphatidylserine (PS), phosphatidylethanolamine (PE), lysophosphatidylethanolamine (LysoPE), phosphatidylcholine (PC) and lysophosphatidylcholine (LysoPC) between *rpl40aΔ* and wild-type (WT) cells. (*Lower panel*) Volcano plots of mitochondria lipidomics presenting the $\log_2$ fold change of abundance of all quantified molecular species of diglycerides, cardiolipin, PG, PA, PI, PS, PE, LysoPE, PC and LysoPC between *rpl40bΔ* and WT cells; $n = 4$.

**Supplementary Table 2.** Quantification of fatty acid content in mitochondria and culture medium using gas chromatography

| | mitochondria | | | | | | culture medium | | | | | |
|---|---|---|---|---|---|---|---|---|---|---|---|---|
| | WT | | *rpl40a*Δ | | *rpl40b*Δ | | WT | | *rpl40a*Δ | | *rpl40b*Δ | |
| Fatty acid | mean [mg/l] | SD | mean [mg/l] | SD | mean [mg/l] | SD | mean [mg/l] | SD | mean [mg/l] | SD | mean [mg/l] | SD |
| **C8:0** | **$0.1^{E}$** | 0.0 | **$0.0^{F}$** | 0.0 | **$0.0^{F}$** | 0.0 | 6.5 | 4.6 | 0.0 | 0.0 | 20.0 | 15.5 |
| **C10:0** | 16.4 | 21.8 | 1.1 | 0.1 | 1.0 | 0.1 | 3.8 | 3.0 | **$3.3^{A}$** | 1.2 | **$6.7^{B}$** | 0.8 |
| C12:0 | 0.7 | 0.2 | 0.2 | 0.0 | 0.4 | 0.2 | 8.4 | 9.2 | 0.5 | 0.2 | 1.7 | 1.2 |
| **C14:0** | **$3.0^{E}$** | 0.4 | **$0.0^{F}$** | 0.0 | **$0.0^{F}$** | 0.0 | 0.0 | 0.0 | 0.0 | 0.0 | 0.0 | 0.0 |
| C16:0 | 5446.2 | 1004.1 | 6314.6 | 292.3 | 5607.9 | 1615.3 | 20.7 | 17.2 | 41.6 | 11.6 | 82.2 | 28.6 |
| C16:1 | 3.8 | 2.2 | 1.3 | 0.3 | 0.8 | 0.3 | 3.7 | 2.3 | 1.8 | 1.5 | 0.8 | 0.2 |
| **C18:0** | 104.7 | 2.4 | 102.6 | 3.6 | 104.5 | 0.9 | **$108.1^{A}$** | 2.7 | **$107.5^{A}$** | 2.6 | **$100.3^{B}$** | 2.5 |
| C18:1 | 4.3 | 1.7 | 0.7 | 0.1 | 1.0 | 0.7 | 6.1 | 3.6 | 2.9 | 1.1 | 2.4 | 0.2 |
| **C18:2** | **$1.0^{A}$** | 0.7 | **$3.1^{B}$** | 0.8 | 3.6 | 1.6 | 5.7 | 2.5 | 6.5 | 3.0 | 3.7 | 0.9 |
| **C20:0** | **$7.7^{E}$** | 8.9 | **$0.0^{F}$** | 0.0 | **$0.0^{F}$** | 0.0 | 20.8 | 18.6 | 17.2 | 9.4 | 11.3 | 15.9 |
| C18:3 | 1.8 | 1.3 | 2.5 | 0.3 | 1.9 | 0.6 | 1.1 | 0.6 | 1.4 | 1.1 | 0.6 | 0.1 |
| C22:0 | 0.5 | 0.4 | 0.5 | 0.2 | 0.2 | 0.0 | 0.6 | 0.4 | 0.4 | 0.4 | 1.0 | 0.3 |
| **C22:1** | **$625.2^{E}$** | 146.5 | **$6562.0^{F}$** | 696.4 | 4829.1 | 3313.4 | 9.3 | 4.8 | 28.5 | 16.0 | 5.6 | 0.9 |
| **C24:0** | **$1543.9^{A}$** | 477.9 | **$299.0^{B}$** | 102.7 | **$490.9^{B}$** | 90.0 | 282.1 | 66.4 | **$299.0^{C}$** | 28.1 | **$150.2^{D}$** | 20.9 |
| UFA | **$636.0^{E}$** | 149.5 | **$6569.7^{F}$** | 696.5 | 4836.4 | 3312.6 | 25.9 | 13.4 | 41.2 | 21.4 | 13.2 | 2.3 |
| SFA | 7123.2 | 552.5 | 6718.0 | 397.4 | 6204.9 | 1705.8 | 450.9 | 81.3 | 469.7 | 15.0 | 373.4 | 50.8 |

The mean and standard deviations of three biological replicates per group. If normally distributed data - Two-sample t test (in the case of not equal variance, then with Welch Correction). In case of not normally distributed data: Kruskal-Wallis ANOVA and Conover's test. Significance: AB – $p \leq 0.05$; CD - $p \leq 0.01$; EF - $p \leq 0.001$. UFA – the sum of all unsaturated fatty acids quantified in the analysis; SFA – the sum of all saturated fatty acids quantified in the analysis. WT, wild-type cells.

**Supplementary Table 4.** Yeast strains used in this study.

| Strain | Genotype | Reference |
|---|---|---|
| **BY4741 (WT)** | *MATa*; his3Δ1; leu2Δ0; met15Δ0; ura3Δ0 | Yeast knockout collection (Horizon Discovery) |
| ***rpl40a*Δ** | YIL148w::kanMX4 | |
| ***rpl40b*Δ** | YKR094c::kanMX4 | |
| ***puf3*Δ** | YLL013c::kanMX4 | |
| **yMS032** | *PUF3*::3xHA | This study |
| **yMS016** | YIL148w::URA3;*PUF3*::3xHA | |
| **yMS014** | YKR094c::URA3;*PUF3*::3xHA | |

**Supplementary Table 5.** The sequence of sense and antisense primers used in this study.

| Gene | Forward (5'-3') | Reverse (5'-3') | Reference |
|---|---|---|---|
| **Primers used for RT-qPCR** | | | |
| *RPL40A* | TGTTTGCCGTAAGTGTTATGCT | GGACGCAATTGGTTGGTGTG | This study |
| *RPL40B* | CCAAAAGGAATCCACTTTACATTTG | CAGTTGTATTTGGAAGCCAAG | |
| *MIA40* | ATGGGGATTCTTTCTATGGC | CACCACTGGAAAGCTCTTC | |
| *PAM16* | CATGGACAAGATTAATAACAGG | GAGCCAGTTCCCATTTTAAC | |
| *TOM20* | ACTCATGCTAAGGAAGTGAAG | CCATTTTCTACGTTGGTGGT | |
| *QCR6* | ACATTTCAAGAACACGGAGG | ACAGTCCTCCTTGTGTTCA | |
| *CCP1* | ACAACGAACAGTGGGACTCT | ACTTGTCCTGGTCATTAGCGT | 1 |
| *TRR1* | CCGTCCCCATTTTCAGAAACA | GCACGCAAATGGTCTTTTCTG | |
| *ACT1* | CATGTTCCCAGGTATTGCCG | GTCAAAGAAGCCAAGATAGA | |
| *ALG9* | TCCATGATACAGGAGCAAGC | CTACCATCAGAACCGCATTC | |
| *TDH1* | GGTATGGCTTTCAGAGTCCCA | AGACAACGGCATCTTCGGTG | |
| **Primers used for quantification of mtDNA copy number** | | | |
| *ACT1* | CACCCTGTTCTTTTGACTGA | CGTAGAAGGCTGGAACGTTG | 2 |
| *MIP1* | CCATCACAAGCAAGAACGGC | GTCCCTTTCCAGCTCAACCA | |
| *MRX6* | CATCCGACGTGGTGCTCTTA | TCTCATCTCTCCCTCCACCC | |
| *COX2* | GTTGATGCTACTCCTGGTAGATT | TTGCATGACCTGTCCCACAC | |
| *COX3* | TTGAAGCTGTACAACCTACC | CCTGCGATTAAGGCATGATG | |
| ***Primers used for yeast strain construction*** | | | |
| *PUF3* | TAAACAAATCTAACTCCCTAGGAAATAG<br>ACATTTAGCCAGTGTTGAGAAACTTGC<br>AGCATTGGTTGAAAATGCGGAGGTGG<br>GAATCTTTTACCCATACGA | ACAAATGGAGGAAAAAAAAAAAAAAA<br>AAAAAAAAAAAAAAATAGTAAAAAGT<br>GAAAGGAGAACGATGATAACACTAAT<br>CATTATTCCTTTGCCCTCGGAC | This study |
| *RPL40A* | *AAATTAAAGGTAGTTGAATCTCTATTTG*<br>*TTGTTGTTATTACCGCTTATTATCCCAT*<br>*AGTTGAGACGACCAAGATTCAAACTAA*<br>*CTATGCGGCATCAGAGC* | TTTAACAGTAGGTATGATAAAAGAAG<br>ATGTACATTAAAGTATACAGTAAATAA<br>ATGTATAGATTGATTGGGCGAAACAG<br>ACCTGATGCGGTATTTTCTCC | |
| *RPL40B* | *ATCTCAAATACACAAATAAGAGCAACCT*<br>*TATATATCACTTTTTCCCGTTCAGCAAG*<br>*AGGTAAAGCCACCAAAGGTTCAAATAA*<br>*CTATGCGGCATCAGAGC* | AATAGAAAAGGAAATATATATGTAGT<br>AAAAAAATCGTGATGTATACAAACTTT<br>GGATTTGTGGAGATCGTAATAAATCG<br>ACCTGATGCGGTATTTTCTCC | |

**Supplementary Table 6.** Parameters for processing and analysis of lipidomics data.

1. Unaligned CD workflow

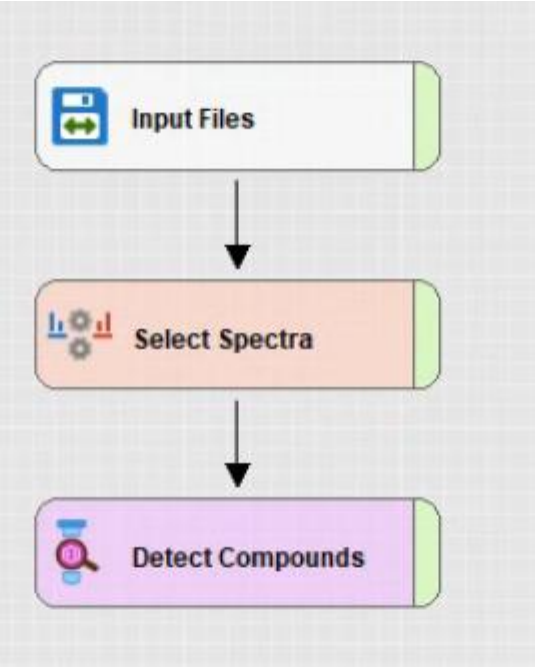


*Figure 1 Graphical representation of Unaligned CD workflow*

| Unaligned Workflow Tree | Workflow Node | Parameters | Value |
|---|---|---|---|
| | **Input files** | | |
| | | NA | NA |
| | **Select Spectra** | | |
| | | Lower RT limit | 0.2 |
| | | Upper RT limit | 30 |
| | | Min. precursor mass [Da] | 100 Da |
| | | Max precursor mass [Da] | 5000 Da |
| | | Polarity mode | Any |
| | | Peak Filters (S/N Threshold) | 1.5 |
| | | Precursor Selection | Use MS(n-1) Precursor |
| | **Detect Compounds** | | |
| | | Mass tolerance [ppm] | 7.5 |
| | | Min. Peak Intensity | 10000 |
| | | Precursor Mass Tolerance [Da] | 0.01 |
| | | Chromatographic S/N Threshold | 1.5 |
| | | Gap Ratio Threshold | 0.35 |
| | | Max. Peak Width [min] | 1 |
| | | Min. Relative Valley | 0.1 |
| | | Compound Assembly (Ions) | [M+H]+1; [M-H+TFA]-1 |
| | | Compound Assembly (Base Ions) | [M+H]+1; [M-H]-1 |

2. Aligned CD workflow

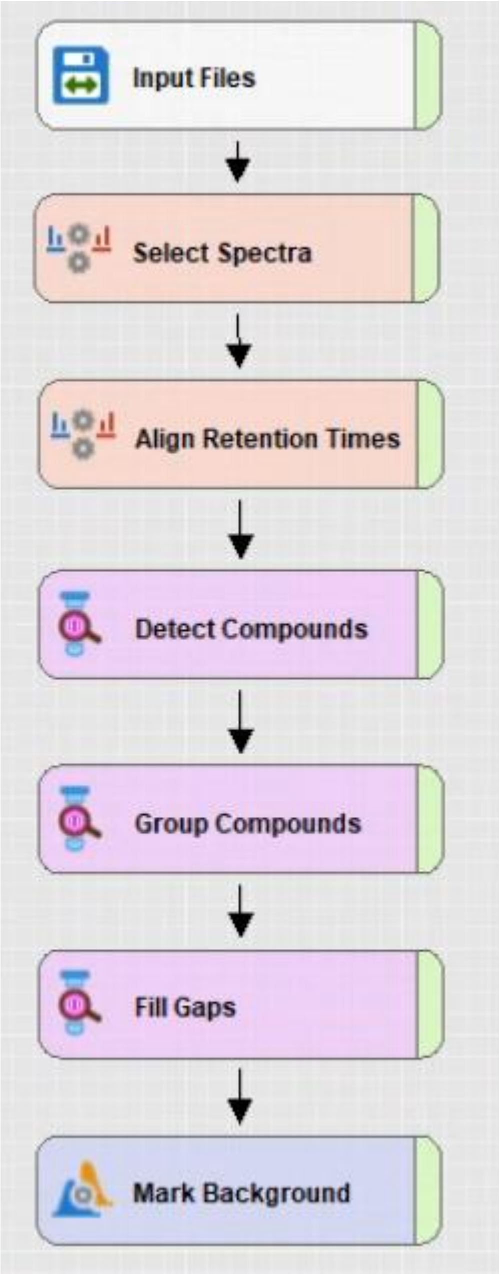


*Figure 2 Graphical representation of Aligned CD workflow*

| Aligned Workflow Tree | Workflow Node | Parameters | Value |
|---|---|---|---|
| | **Input files** | | |
| | | NA | NA |
| | **Select Spectra** | | |
| | | Lower RT limit | 0.2 |
| | | Upper RT limit | 30 |
| | | Min. precursor mass | 100 Da |
| | | Max precursor mass | 5000 Da |
| | | Polarity mode | Any |
| | | Peak Filters (S/N Threshold) | 1.5 |
| | | Precursor Selection | Use MS(n-1) Precursor |
| | **Align Retention Times** | | |
| | | Alignment model | Adaptive curve |
| | | Alignment Fallback | Use Linear Model |
| | | Maximum shift [min] | 0.6 |
| | | Mass tolerance [ppm] | 7.5 |
| | | Remove outlier | True |
| | **Detect compounds** | | |
| | | Mass tolerance [ppm] | 7.5 |
| | | Min. Peak Intensity | 10000 |
| | | Precursor Mass Tolerance [Da] | 0.025 |
| | | Chromatographic S/N Threshold | 1.5 |
| | | Gap Ratio Threshold | 0.35 |
| | | Max. Peak Width [min] | 1 |
| | | Min. Relative Valley | 0.1 |
| | | Compound Assembly (Ions) | [M+H]+1; [M-H+TFA]-1 |
| | | Compound Assembly (Base Ions) | [M+H]+1; [M-H]-1 |
| | **Group compounds** | | |
| | | Mass tolerance [ppm] | 7.5 |
| | | RT tolerance [min] | 0.2 |
| | | Minimum valley [%] | 10 |
| | | Align peaks | False |
| | | Preffered ions | [M+H]+1; [M-H+TFA]-1 |
| | | Area Integration | Most Common Ion |
| | | Peak Rating Threshold | 0 |
| | **Fill gaps** | | |
| | | Mass tolerance [ppm] | 7.5 |
| | | S/N threshold | 1.5 |
| | | Use Real Peak Detection | True |
| | | Apply Restrictive Gap Filling | True |
| | | Min # Scans per Peak | 3 |
| | **Mark Background** | | |
| | | Max. Sample/Blank | 3 |
| | | Max. Blank/Sample | 0 |

| | |
|---|---|
| Hide Backgroud | True |

### 3. Lipidex analysis

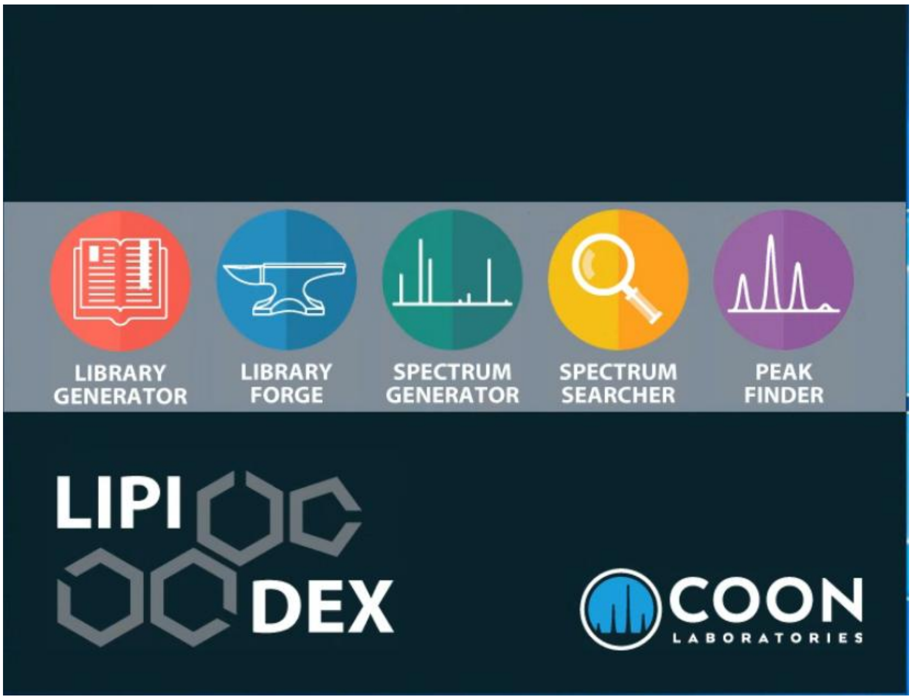


*Figure 3 Central screen of the Lipidex 1.1.0 version with the key elements of its algorithm*

#### 3.1. Spectrum Searcher

*Table 1 Summary of Spectrum Searcher algorithm parameters*

| | |
|---|---|
| MS1 Search Tolerance (Th) | 0.01 |
| MS2 Search Tolerance (Th) | 0.01 |
| Mas Search Results Returned | 1 |
| MS2 Low Mass Cutoff (Th) | 61.00 |

#### 3.2. Peak Finder

| | |
|---|---|
| Min. Lipid Spectral Purity (%) | 75 |
| Min. MS2 Search Dot Product | 500 |
| Min. MS2 Search Rev. Dot Product | 700 |
| FWHM Window Multiplier | 2.0 |
| Mas. Mass Difference (ppm) | 15 |

**Supplementary Table 8. Summary of chromatographic binary gradient steps.** The total method run time was 35 min.

| Time [min] | Mobile phase A [%] | Mobile phase B [%] |
|---|---|---|
| ***Initial*** | 98 | 2 |
| **2.0** | 70 | 30 |
| **5.0** | 50 | 50 |
| **6.0** | 15 | 85 |
| **20.0** | 15 | 85 |
| **21.0** | 5 | 95 |
| **28.0** | 5 | 95 |
| **29.0** | 98 | 2 |

#### Supplementary References


1. Gościńska, K., Shahmoradi Ghahe, S., Domogała, S., and Topf, U. (2020). Eukaryotic Elongation Factor 3 Protects Saccharomyces cerevisiae Yeast from Oxidative Stress. Genes (Basel) *11*.
2. Seel, A., Padovani, F., Mayer, M., Finster, A., Bureik, D., Thoma, F., Osman, C., Klecker, T., and Schmoller, K.M. (2023). Regulation with cell size ensures mitochondrial DNA homeostasis during cell growth. Nat Struct Mol Biol *30*, 1549-1560.
